# PBRM1 directs PBAF to pericentromeres and protects centromere integrity

**DOI:** 10.1101/2024.06.29.601326

**Authors:** Karen A. Lane, Alison Harrod, Lillian Wu, Theodoros I. Roumeliotis, Hugang Feng, Shane Foo, Katheryn A. G. Begg, Federica Schiavoni, Frank T. Zenke, Alan A. Melcher, Jyoti S. Choudhary, Jessica A. Downs

## Abstract

The specialised structure of the centromere is critical for effective chromosome segregation, but its repetitive nature makes it vulnerable to rearrangements.

Centromere fragility can drive tumorigenesis, but protective mechanisms preventing fragility are still not fully understood. The PBAF chromatin remodelling complex is frequently misregulated in cancer, but its role in cancer is still not fully characterized. Here, we identify PBAF as a protector of centromere and pericentromere structure with profound consequences for genome stability. A conserved feature of isogenic cell lines lacking PBRM1, a subunit of PBAF, is compromised centromere and pericentromere integrity. PBAF is present at these regions, and the binding pattern changes when PBRM1 is absent. PBRM1 loss creates a dependence on the spindle assembly checkpoint, which represents a therapeutic vulnerability. Importantly, we find that even in the absence of any perturbations, PBRM1 loss leads to centromere fragility, thus identifying a new player in centromere protection.

## Introduction

The centromere is a specialised region of the chromosome that is responsible for the attachment of spindle fibres via the kinetochore during mitosis, ensuring that each daughter cell receives an equal and identical set of chromosomes. Kinetochore assembly is a highly regulated process that builds upon the constitutive centromere associated network (CCAN). The CCAN bridges the CENPA-containing centromeric chromatin with the outer kinetochore components. CENPB contributes to this assembly by bridging CENPA and the CCAN subunit CENPC. Human centromeric DNA is made up of repetitive arrays of alpha satellite (α-Sat) repeats forming higher order repeats (HORs)^1^. Many α -Sat repeats contain a motif termed the B-box, which is bound by CENPB. CENPB is important for creating three dimensional structures in the centromere that promote appropriate chromosome segregation^2^.

Centromeres are flanked by pericentromeres or transition regions, which are made up of a wider range of repetitive sequences in human cells, including α -Sat and non- α -Sat repeats, transposable elements, and duplications^1^. H3K9me3 and other repressive marks are enriched in pericentromeres and the repetitive elements are mostly silenced^3^. However, pericentromeric regions also contain non-repetitive elements, including protein coding genes, some of which are expressed^1^. Therefore, while largely heterochromatic, there are interspersed regions of open chromatin in pericentromeric regions and transition arms.

The repetitive nature of centromeric and pericentromeric sequences facilitates topological organisation, but this comes at a cost, since these repetitive sequences are vulnerable to inappropriate rearrangements^4, 5^. Moreover, recent work demonstrated that cells use centromere-associated DNA breaks to help specify functional centromeres^6^, which increases vulnerability if these are processed or repaired inappropriately. Pathological centromere fragility contributes to human disease, such as cancer^4^. Cells must therefore balance the use of these specialised and repetitive structures with mechanisms that protect genome integrity. How this is achieved is not yet fully understood.

PBRM1 (or BAF180) is a subunit of the PBAF chromatin remodelling complex, one of three mammalian SWI/SNF remodelling complexes. PBRM1 is frequently mutated in cancer, and evidence support the idea that PBRM1 can function as a tumour suppressor^7^. Loss of function mutations are particularly prevalent in clear cell renal cell carcinoma (ccRCC), but they are also found across a range of other cancer types^8^. A critical question is what the fundamental functions of PBRM1 are, which, when lost, contribute to the development or evolution of cancer.

Here, we identify PBRM1 as a factor that prevents centromere fragility. We show that cells lacking PBRM1 have lower levels of centromere- and pericentromere-associated proteins and have altered patterns of organisation of these structures in cells. PBRM1 and the SMARCA4 subunit of PBAF are physically present at these regions, and the SMARCA4 binding pattern changes when PBRM1 is absent. Furthermore, PBRM1 loss leads to mitotic defects and creates a dependence on the spindle assembly checkpoint, which represents a therapeutic vulnerability. Importantly, we find that even in the absence of any perturbations, PBRM1 loss leads to centromere fragility, thus identifying a new player in centromere protection.

## Results

### Analysis of isogenic PBRM1 knockout (KO) cell lines identifies misregulation of centromere- and pericentromere-associated proteins

To identify core functions of PBRM1, we generated a panel of 17 clonal cell lines with CRISPR- Cas9 engineered loss of function mutations in PBRM1 across five different cell line backgrounds, including both cancer-derived and immortalised non-cancerous parental cell lines (Fig. 1a and ^9^). The growth rate, cell cycle profile, and morphological changes in the knockout (KO) cells were analysed relative to the parental lines (Fig. 1b-f, and Supplementary Fig. 1). We found none of these features were substantially altered in any of the cell lines other than a modestly reduced growth rate when PBRM1 is lost (Fig. 1d and Supplementary Fig. 1).

**Figure 1.**
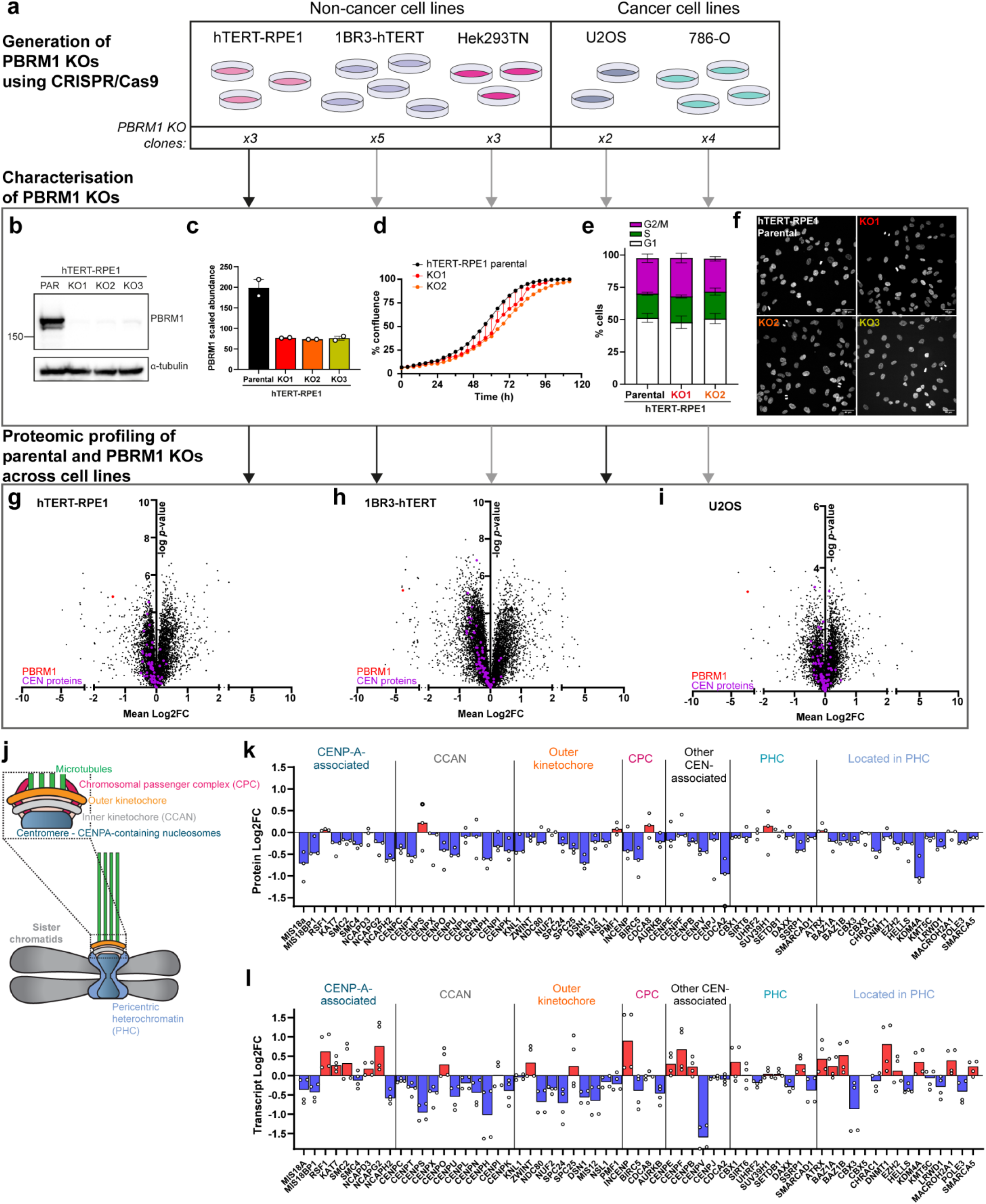
Analysis of isogenic PBRM1 knockout (KO) cell lines identifies centromere associated protein misregulation as a common feature. **(a)** Workflow for generation of PBRM1 knockouts in a panel of cell lines. Number of independent clones validated for each cell line is indicated at the bottom of the schematic. (b)-(f) Characterisation of PBRM1 knockouts using knockouts in the hTERT-RPE1 cell line as an example. (b) Western blotting of whole cell lysates from parental and PBRM1 knockout cells for PBRM1. α-tubulin is used as a loading control. (c) Scaled abundances of PBRM1 in proteomic analyses of whole cell protein extracts. Points correspond to independent biological replicates (*n*=2). (d) Proliferation of RPE1 parental and two PBRM1 knockout clones, measured using phase contrast Incucyte images (*n*=2). (e) Cell cycle distribution of RPE1 parental and two PBRM1 KO clones measured using flow cytometry. *n*=4, mean±SEM, and data were non-significant (ns) based on a 2way ANOVA using Dunnett’s multiple comparisons test. (f) Immunofluorescence images of nuclear morphology in RPE1 parental and PBRM1 knockout clones. Scale bar corresponds to 40µm. (g)-(i) Protein abundances in PBRM1 knockouts compared to parental cells in (g) hTERT-RPE1, (h) 1BR3-hTERT, and (i) U2OS cell lines, detected using LC-MS of whole cell protein extracts. The mean log2 fold change (Log2FC) of protein abundance in PBRM1 knockouts versus parental cells is plotted against the -log *p* value. PBRM1 is highlighted in red, while centromere- & pericentromere-associated proteins are highlighted in purple. (j) Schematic outlining regions of the centromere and pericentric heterochromatin, including the kinetochore in mitosis. (k) Median Log2FC of annotated centromere- & pericentromere-associated proteins in RPE1 PBRM1 knockouts compared to parental cells. Points correspond to individual knockout clones from one of two independent biological replicates. (l) Transcript levels of annotated centromere- & pericentromere-associated genes corresponding to the proteins in (k) were detected using RNA-seq. Median Log2FC of annotated genes transcribing centromere- & pericentromere-associated proteins in RPE1 PBRM1 knockouts was plotted compared to parental cells. Points correspond to individual knockout clones from two independent biological replicates.

To characterise the molecular profiles of the cells, we performed mass spectrometry on all PBRM1 KO clones and the corresponding parental lines. Pathway analysis showed alterations in chromatin organisation, DNA repair and recombination, and innate immune signalling were apparent across cell lines (Supplementary Fig. 2a and 2b). Previously, we found that PBRM1 is important for mediating sister chromatid cohesion at centromeres^10^, raising the possibility that PBRM1 is important for chromatin structure and organisation at or near centromeres. We therefore interrogated the proteomic datasets for centromere- and pericentromere-associated proteins, including CENPA interacting proteins, the constitutive centromere-associated network (CCAN) complex, the outer kinetochore, the chromosomal passenger complex (CPC), pericentromeric heterochromatin proteins, and other annotated peri/centromere associated proteins. Individual protein levels were modestly altered in the PBRM1 KO cells, but looking across the pathways, a trend was apparent in the PBRM1 KO cells when compared with their isogenic parental cell lines (Fig. 1g-k, and Supplementary Fig.2c,d and 3a). In contrast, transcript levels of these genes were not consistently downregulated in the PBRM1 KO cells, indicating that the protein level changes were not primarily being driven by misregulation of gene expression (Fig. 1l, and Supplementary Fig. 3b,c). Moreover, downregulation was specific to centromere-associated complexes; by contrast, no consistent changes were apparent when centrosome proteins were analysed (Supplementary Fig. S3e,f).

We further investigated available proteomic datasets in the cancer cell line encyclopedia (CCLE) to explore whether downregulated centromere proteins are a general feature of PBRM1 loss. In the absence of isogenic comparisons, we ranked cell lines according to PBRM1 protein levels. Notably, we found a correlation between PBRM1 protein levels and peri/centromere-associated proteins (Supplementary Fig. 3d). These data suggest that lower levels of centromere and pericentromere-associated proteins is a common feature of cells lacking PBRM1 expression.

### Loss of PBRM1 results in altered organization of centromeres and pericentromeric heterochromatin

We next set out to understand whether the decreased level of centromere and pericentromere-associated proteins had any functional consequence on their organisation. We first looked to see whether there were any detectable changes in centromere structure using FISH probes against centromere α-Sat repeats in chromosome 2 or 10. We found that the area of the signal in the KO clones was greater than that of the parental RPE1 or 1BR3 cells (Fig. 2a-c and Supplementary Fig. 4a-g). One possible explanation for the changes in FISH signals is a failure to form appropriate three-dimensional structures in the KO cells, leading to an increased volume of α-Sat repeat-containing chromatin in the KO cell nuclei. This pattern was previously observed in CENPA-depleted cells^11^ raising the possibility that PBRM1 loss leads to CENPA deficiency. Since CENPA was not present in our proteomic dataset, we interrogated CENPA levels by IF. However, we found no clear difference in CENPA levels in the PBRM1 KO cells (Supplementary Fig. 4h-j).

**Figure 2.**
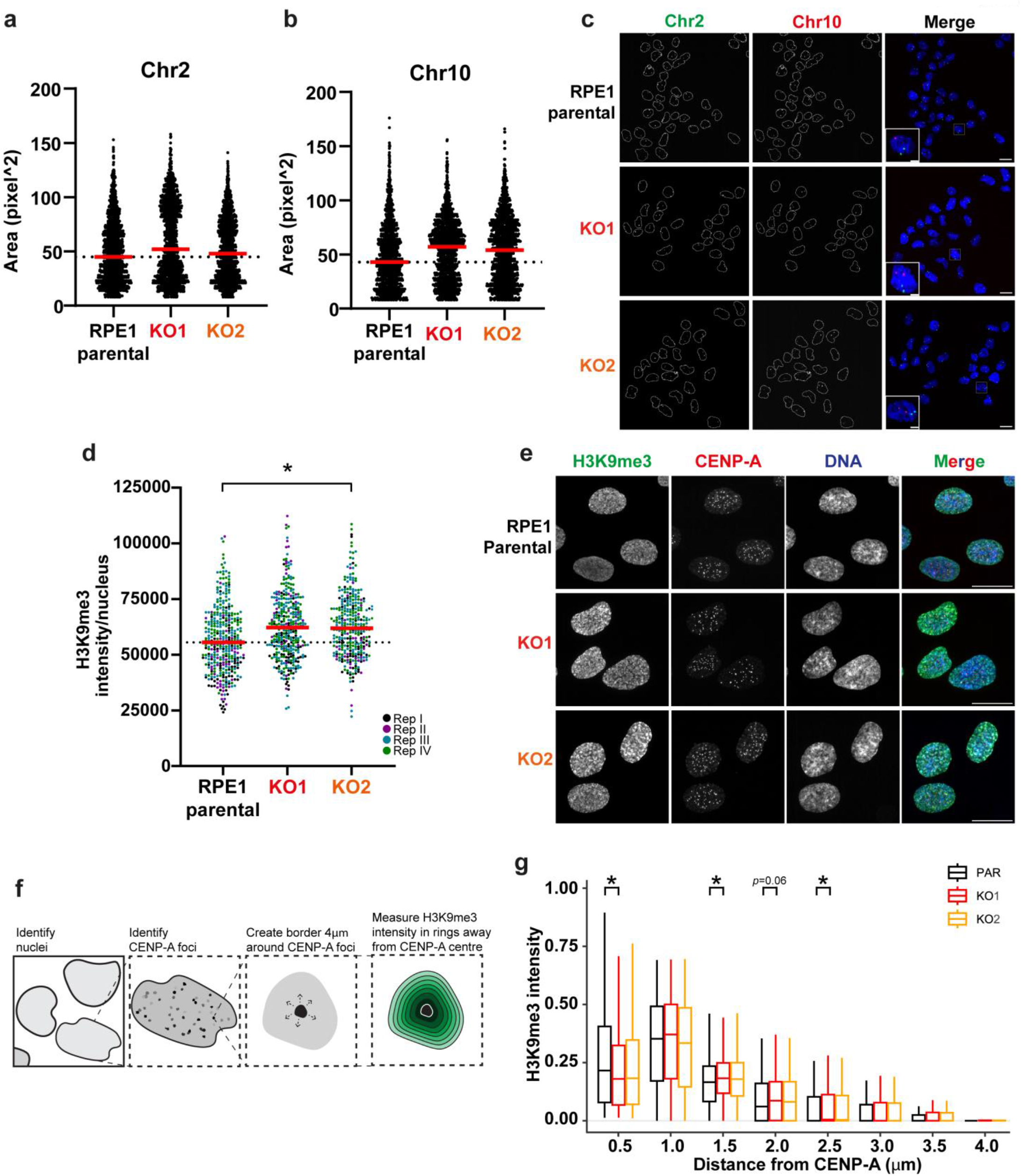
PBRM1 KO cells display increased H3K9me3 intensity around centromeres. **(a)-(b)** Quantification of the area of individual foci in RPE1 parental or PBRM1 knockout cells stained for α-satellite centromeric regions in (a) chromosome 2 and (b) chromosome 10, using FISH probes against α-satellite sequences in the corresponding chromosomes. *n*=3, line at median. **(c)** Representative images of α-satellite FISH of chromosomes 2 and 10 in RPE1 parental and PBRM1 knockouts. Scale bars corresponds to 20µm; in zoomed inset images, scale bars correspond to 5µm. **(d)** Quantification of total H3K9me3 signal per nucleus in RPE1 parental and PBRM1 KOs. Points correspond to individual nuclei. *n*=4, with red line indicating the median intensity and dotted line corresponding to the median intensity of parental cells. At least 325 nuclei were analysed per condition and data were analysed using t-test of experiment medians, \**p*<0.05. **(e)** Representative images showing H3K9me3 and CENP-A signal in RPE1 parental and PBRM1 KO cells. Scale bars correspond to 20μm. **(f)** Schematic describing method of quantifying H3K9me3 signal around centromeres, which were defined the presence of CENP-A. Briefly, a Cell Profiler pipeline was used to first identify nuclei and CENP-A foci within each nucleus. CENP-A signal was then expanded to a total distance of 4µm from the centre of each CENP-A focus, divided into 8 rings. H3K9me3 intensity was then measured in each of these rings. **(g)** Boxplot indicating quantifying H3K9me3 intensity at increasing distances from the centre of CENP-A foci in RPE1 parental and PBRM1 KOs. Boxes show a line at median and corresponding quartiles, *n*=3. CENP-A foci in at least 290 nuclei quantified per condition and data were analysed using t-test of experiment medians, \**p*<0.05.

We next looked to see whether there were any detectable changes in pericentromeric heterochromatin. We performed immunofluorescence using an antibody against H3K9me3, a marker of heterochromatin, and found that the intensity of this signal is subtly but consistently increased in the absence of PBRM1 (Fig. 2d,e), suggesting altered organisation or prevalence of heterochromatin in these cells. To see whether there were changes specifically in pericentromeric heterochromatin, we used an antibody against CENPA to identify centromeres and then quantified the surrounding H3K9me3 signal (Fig. 2f). Interestingly, we find that the pattern of H3K9me3 intensity changes in both PBRM1 KO clones (Fig. 2g). In both cases, the signal is lower in the shell closest to CENPA, but higher in outer shells, suggesting that PBRM1 helps to organise chromatin in regions surrounding centromeres.

### PBRM1 and SMARCA4 are present at pericentromeres, and PBRM1 loss leads to changes in SMARCA4 association

One possible mechanism by which PBRM1 is regulating the structure and organisation of chromatin and associated proteins in centromeres and pericentromeric regions is through direct remodelling activity in these regions. There is some evidence to support this possibility. PBAF was reported to associate with kinetochores of mitotic chromosomes^12^, and SWI/SNF subunits have been identified in protein-interaction studies of centromere-associated proteins, including CENPC, INCENP, and Bub1^13, 14, 15^. However, in contrast to our understanding of PBAF binding patterns in euchromatic regions of the genome, little is known about the specific binding pattern of PBAF in repetitive regions.

We therefore set out to determine whether PBAF associates with centromeric or pericentromeric chromatin (including HORs, HSAT repeats, transition (ct) arms; Fig. 3a) and gain a comprehensive view of PBAF localisation patterns. To do this, we performed CUT&RUN sequencing in both low and high salt conditions to ensure capture of heterochromatic regions^16^. We mapped PBRM1 and SMARCA4 (BRG1), one of two catalytic subunits of PBAF, using both IgG and a SMARCA4 KO cell line (Fig. 3a and Supplementary Fig. 6b) as negative controls, and CENPB as a positive control. Because of the repetitive nature of the region, we analysed the data in two ways: peak calling of uniquely mapping reads, and a *k*-mer analysis of reads mapping to these regions, thus allowing analysis of reads mapping to multiple locations (Fig. 3a and Supplementary Fig. 5, described below).

**Figure 3.**
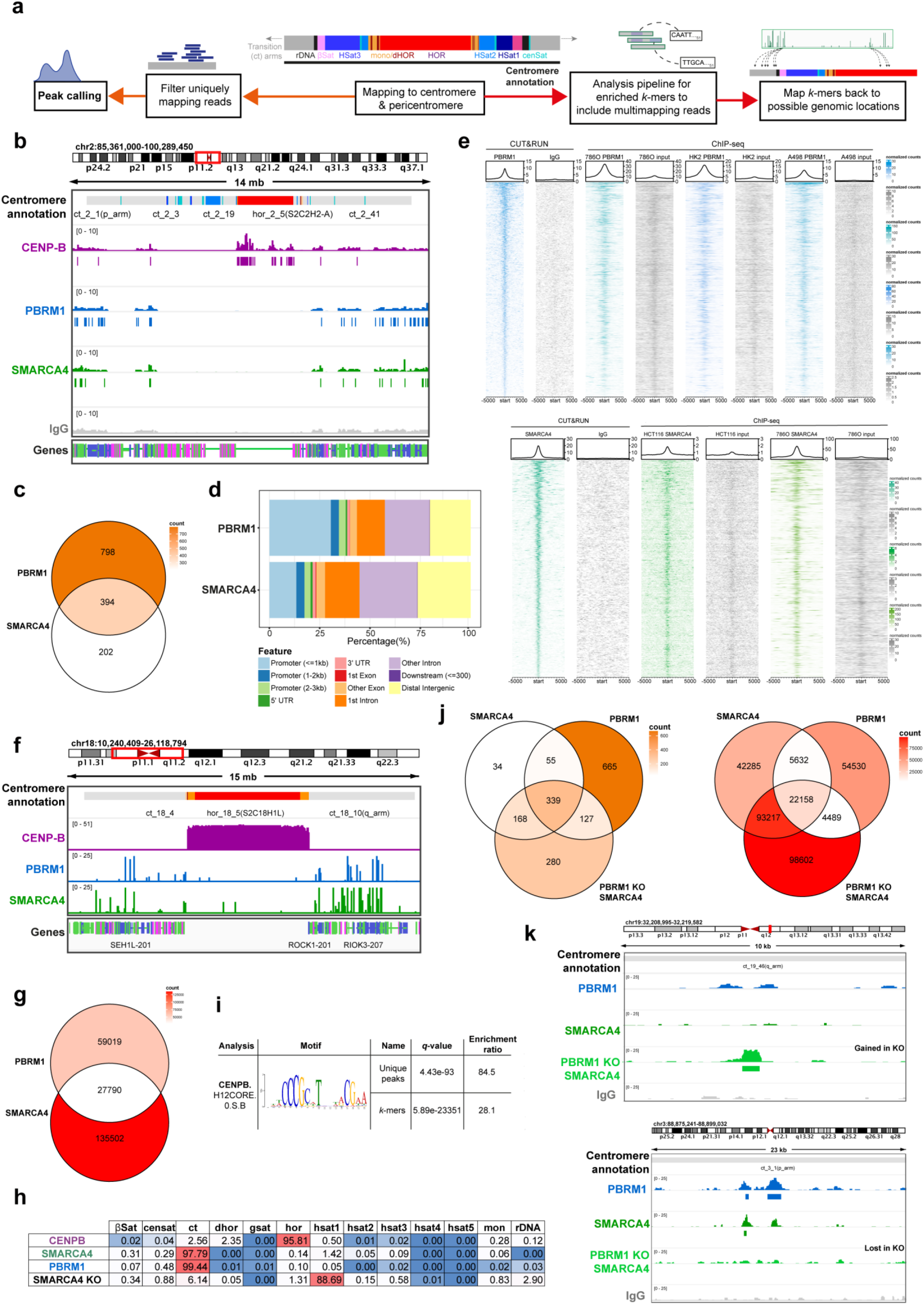
SMARCA4 is present at centromeres, and this pattern changes in PBRM1 KOs. **(a)** Simplified flowchart describing the mapping strategy to centromeric and pericentromeric sequences (more detailed version in Fig S5). **(b)** Representative genome tracks displaying coverage of reads from CENP-B (purple), PBRM1 (blue), SMARCA4 (green) and IgG control (grey) CUT&RUN sequencing in RPE1 parental cells across the centromere. **(c)** Venn diagram indicating the overlap of significantly enriched SMARCA4 and PBRM1 peaks in centromeric and pericentromeric regions in RPE1 parental cells. **(d)** Stacked colour bar representing the genomic distribution of enriched PBRM1 and SMARCA4 peaks, categorised by feature, within centromeric and pericentromeric regions. **(e)** CUT&RUN and ChIP-seq signal heatmaps (PBRM1 – top, blue; SMARCA4 – bottom, green) in the indicated cell lines +/-5kb from the centre of RPE1 PBRM1 or SMARCA4 CUT&RUN peaks in the centromere and pericentromere, versus the IgG or input control (grey) of each experiment. Peaks are ordered by signal of the left-most heatmap, i.e. CUT&RUN peaks. An average signal plot is shown at the top of each heatmap. **(f)** Representative genome tracks displaying mapping locations of enriched *k*-mers across the centromere and pericentromere from analysis of CENP-B (purple), PBRM1 (blue) and SMARCA4 (green) CUT&RUN sequencing in RPE1 parental cells. **(g)** Venn diagram indicating the overlap of enriched *k*-mers (FC>2) in centromeric and pericentromeric regions, in SMARCA4 and PBRM1 in RPE1 parental cells. **(h)** Percentage of enriched *k*-mers that map to specific regions in the centromere and pericentromere in each condition, including negative control of SMARCA4 *k*-mers enriched in SMARCA4 KO cells. **(i)** CENP-B motif detection following motif analysis of CENP-B-enriched *k*-mers compared to a shuffled control, where CENP-B *k*-mer sequences were shuffled maintaining 3-mer frequencies. **(j)** Venn diagram indicating the overlap of significantly enriched peaks (left, orange) or *k*-mers (right, red) in the centromere and pericentromere, in SMARCA4 and PBRM1 in RPE1 parental cells, and SMARCA4 in PBRM1 knockout (KO) cells. **(k)** Representative genome tracks displaying coverage of reads from PBRM1 (blue), SMARCA4 (green) and IgG control (grey) CUT&RUN sequencing in RPE1 parental cells and SMARCA4 in PBRM1 knockout cells (light green), showing an example of peaks gained (top) or lost (bottom) in the PBRM1 knockout cells. For genome browser tracks (b, k), one representative independent biological replicate is shown, with boxes underneath representing peaks that were called as significantly enriched (*q*<0.01) in at least two out of the three replicates versus their IgG control. Genomic location is indicated at the top, centromeric annotation is shown above the tracks and transcript annotation is shown below (b, f, k). Venn diagrams (c, g, j) show enriched peaks or k-mers found in at least two out of the three independent biological replicates, versus their IgG controls. The colour corresponds to the total number of enriched peaks or *k*-mers in each region of the Venn diagram (count).

We first aligned uniquely mapping reads using the reported telomere-to-telomere (T2T) CHM13 genome^1, 17^. CENPB localised primarily to HORs, as expected, with some additional sites of enrichment in flanking regions (Fig. 3b and Supplementary 6a). We found enrichment of SMARCA4 and PBRM1 primarily in pericentromere sequences when compared with negative controls, and most SMARCA4 and PBRM1 peaks were located in the transition (ct) arms (Fig. 3b and Supplementary Fig. 6a). As expected, many of the PBRM1 and SMARCA4 peaks overlapped, but non-overlapping peaks were also identified (Fig. 3c, Supplementary Fig. 6c,e and f). This likely reflects the combinatorial flexibility of the complexes (*i.e.* PBAF can contain SMARCA2 instead of SMARCA4, and SMARCA4 is found in other SWI/SNF complexes).

When analysing the peaks found in centromeres and pericentromeric regions, we find that the peak distribution profile of PBRM1 has a stronger association with promoters when compared with SMARCA4 (Fig. 3d). This profile is similar to that of the genome-wide distributions of both PBRM1 and SMARCA4 in our dataset (Supplementary Fig. 6d), and is consistent with previous studies showing that, in euchromatin, PBRM1-containing PBAF complexes are enriched at both promoters and enhancers, whereas BAF complexes are more often found at enhancers^18, 19^. To determine whether pericentromere-association is a conserved feature of PBAF, we interrogated datasets in which PBRM1 and SMARCA4 were mapped^20, 21^, and strikingly, we find that the association of PBRM1 and SMARCA4 in these regions is apparent in other cell line models (Fig. 3e).

Because of the repetitive nature of these regions, we also analysed centromere and pericentromere-associated reads that mapped to multiple locations, and were therefore unable to be precisely mapped, by performing a *k*-mer analysis^1, 22^ to identify 51-mer sequences that are significantly enriched (Supplementary Fig. 5) in the mapping datasets (see Fig. 3a and Supplementary Fig. 5 for workflow) relative to the IgG control. CENPB was analysed relative to IgG as a positive control.

In order to look for binding patterns and proximity to other features, enriched centromere- and pericentromere-associated *k*-mers were mapped back onto the T2T genome. The CENPB-associated *k*-mers map primarily to the active higher order repeats (HORs) where CENPB is known to bind (Fig. 3f,h, and Supplementary 6g,h), and motif analysis of the CENPB associated *k*-mers identified the CENPB box (Fig. 3i), providing support for the utility of this approach.

Similar to peak enrichment, we found that the majority of PBRM1- and SMARCA4-associated *k*-mers are located within the transition arms, but a small proportion map to repeats (Fig. 3f-h, Supplementary Fig. 6g,h), which may indicate some association across these regions. Again, we find both overlapping as well as PBRM1- and SMARCA4-specific *k*-mers (Fig. 3g and Supplementary Fig. 6i).

We additionally mapped SMARCA4 in PBRM1 KO cells to interrogate changes to PBAF binding patterns when PBRM1 is absent. A subset of SMARCA4-enriched peaks and SMARCA4-enriched *k*-mers are lost when PBRM1 is absent (Fig. 3j,k and Supplementary Fig. 7), suggesting that PBRM1 is important for targeting SMARCA4 and PBAF to these locations. Interestingly, a considerable number of SMARCA4-enriched peaks and SMARCA4-enriched *k*-mers are gained when PBRM1 is deficient (Fig. 3j,k and Supplementary Fig. 7), suggesting that PBRM1 loss leads to aberrant SMARCA4 binding at sites not normally bound by SMARCA4. Together, these data indicate that PBAF binds to specific sites in chromatin flanking centromeres, and this binding pattern is disrupted when PBRM1 is deficient.

### Distinct patterns in PBRM1-dependent and -independent SMARCA4 association with pericentromeric regions

We investigated the relationship between PBRM1 and SMARCA4 peaks and previously mapped histone post-translational modifications (PTMs;^23^). As expected, H3K9me3 is enriched in the transition arms (Fig. 4a and Supplementary Fig. 8a), but not uniformly, and we find that PBRM1 and SMARCA4 are enriched in areas where H3K9me3 is low and PTMs associated with open chromatin, such as H3K4me3 and H3K9ac, are present (Fig. 4a and Supplementary Fig. 8a).

**Figure 4.**
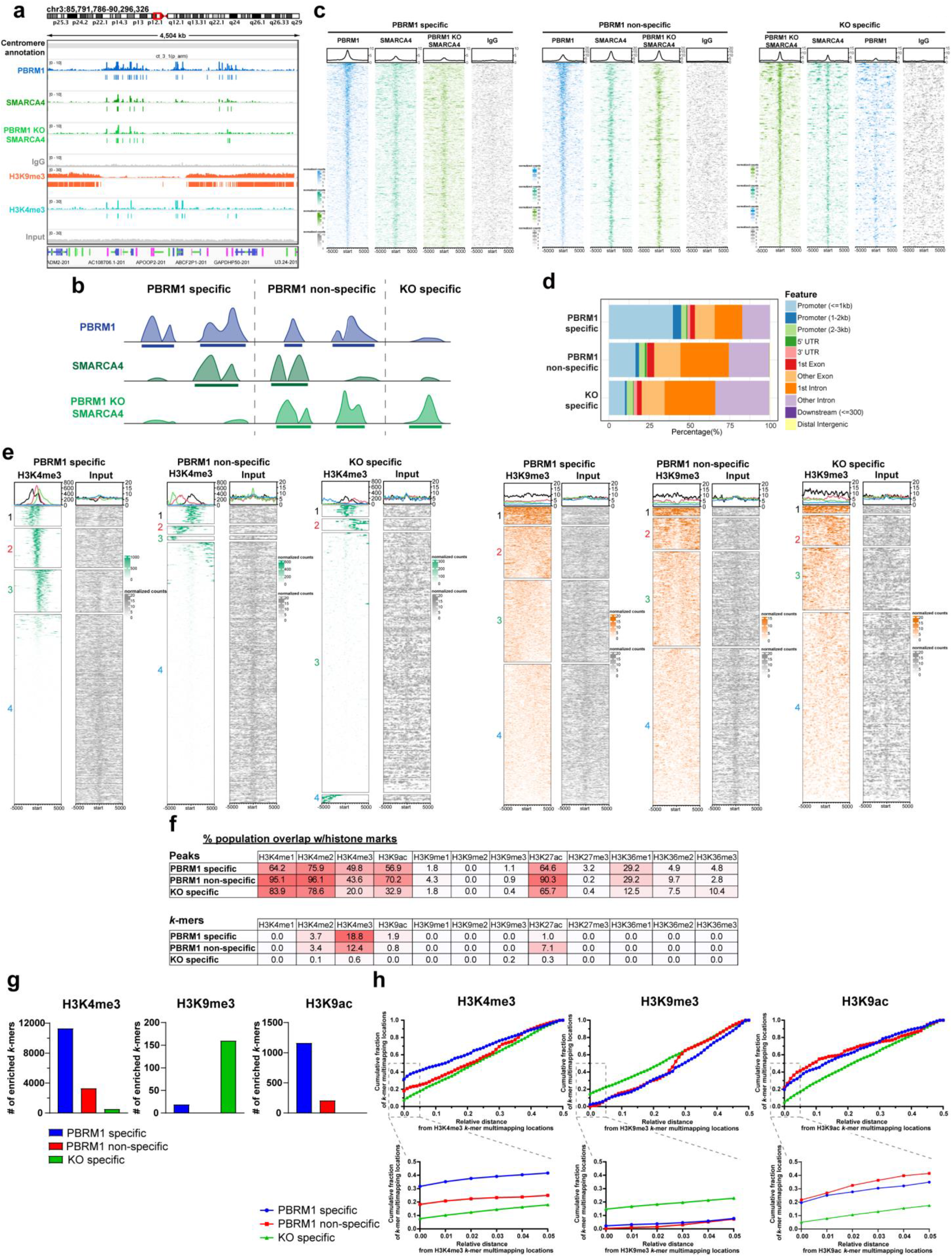
SWI/SNF enrichment at a subset of centromeric chromatin marks is altered in a PBRM1-dependent manner. **(a)** Representative genome tracks displaying coverage of reads from PBRM1 (blue), SMARCA4 (green) and IgG control (grey) CUT&RUN sequencing in RPE1 parental cells, SMARCA4 in PBRM1 knockout cells (light green), and H3K9me3 (orange) and H3K4me3 (teal) RPE1 ChIP-seq showing the boundary of heterochromatin. One representative independent biological replicate is shown, with boxes underneath representing peaks that were called as significantly enriched (*q*<0.01) in at least two replicates versus their IgG or input control. Genomic location is indicated at the top, centromeric annotation is shown above the tracks and transcript annotation is shown below. **(b)** Schematic showing method of identification of PBRM1 specific, PBRM1 non-specific and KO specific subsets of peaks from CUT&RUN sequencing. Example graphics for called peaks are shown for each subset. **(c)** CUT&RUN signal heatmaps representing the three subsets of peaks in the centromere and pericentromere. Signal +/- 5kb from the centre of each peak is shown for PBRM1 (blue) and SMARCA4 in parental RPE1 (green), SMARCA4 in PBRM1 knockout cells (light green) and IgG control (grey). Peaks are ordered by signal of the left-most heatmap for each condition. An average signal plot is shown at the top of each heatmap. **(d)** Stacked colour bar representing the genomic distribution of the three subsets of peaks, categorised by feature, within centromere and pericentromere regions. **(e)** RPE1 ChIP-seq signal heatmaps representing the three subsets of peaks in the centromere and pericentromere. Signal +/- 5kb from the centre of each peak is shown for H3K4me3 (green, left), H3K9me3 (orange, right) and input control (grey). Heatmaps are split into four clusters based on the signal of H3K4me3 (left) or H3K9me3 (right) for each subset of peaks. Average signal plots for each cluster are shown at the top of each heatmap. **(f)** Tables containing the percentage overlap of each subset of peaks (top) or *k*-mers (bottom) from CUT&RUN sequencing data with enriched histone mark peaks or *k*-mers from RPE1 ChIP-seq data, specifically in centromeric and pericentromeric regions. A colour scale of low (white) to high (red) is shown on each table. **(g)** Graphs indicating the number of *k*-mers enriched in subsets (PBRM1 specific (blue), PBRM1 non-specific (red), and KO specific (green)) overlapping with enriched H3K4me3 (left), H3K9me3 (middle), or H3K9ac (right) *k*-mers. **(h)** Plots displaying the cumulative fraction of reference *k*-mer multimapping locations (PBRM1 specific (blue), PBRM1 non-specific (red) and KO specific (green)) at a relative distance from query histone mark *k*-mer multimapping locations (H3K4me3, left; H3K9me3, middle; H3K9ac, right). Relative distances are calculated as the shortest distance between each *k*-mer multimapping location in the query set and the two closest *k*-mer multimapping locations in the reference set, divided by the total distance between the two closest k-mer multimapping locations, so the maximum relative distance is 0.5. A zoomed in view of the smallest relative distances (0 to 0.05) is shown below each plot.

To further explore binding patterns, we divided centromere- and pericentromere-specific sites of PBRM1 and SMARCA4 chromatin association (both peaks and *k*-mers) into three different groups (Fig. 4b,c, and Supplementary Fig. 8b). First, we looked at all PBRM1 enriched sites that did not have SMARCA4 enrichment in the PBRM1 KO cells (PBRM1 specific), representing sites that are dependent on PBRM1. Second, we grouped PBRM1 enriched sites present at locations where SMARCA4 is still bound when PBRM1 is deficient (PBRM1 non-specific), which likely represent other SWI/SNF complexes, such as BAF. Third, we grouped all SMARCA4 enriched sites that appear only when PBRM1 is deficient (KO specific), which reflect potentially aberrant or inappropriate binding when PBRM1 is deficient (Fig. 4b,c and Supplementary Fig.8b-e). As expected, a greater proportion of PBRM1 specific sites of enrichment are located in promoter regions when compared with PBRM1 non-specific sites (Fig. 4d).

We then compared the location of these groups of enriched sites (peaks and *k*-mers) to mapped PTMs. Again, consistent with genome-wide binding patterns of BAF and PBAF, we find that the PBRM1 specific peaks are associated with both promoter- and enhancer-associated PTMs, such as H3K4me3 (Fig. 4e,f), and PBRM1 non-specific SMARCA4 peaks show relatively more association with enhancer-associated PTMs, such as H3K27ac (Fig. 4f).

Interestingly, the SMARCA4 peaks that arise when PBRM1 is deficient (KO specific) display a distinct pattern from either of the other two groups. There is a decrease in association with regions normally enriched in H3K4me3, H3K9ac, and H3K36me1, and an increase in association with H3K36me3-enriched chromatin (Fig. 4f). This could reflect changes in chromatin binding preferences of residual SMARCA4-containing complexes, or changes in the chromatin composition of these regions when PBRM1 is deficient.

In addition, we noticed a small but potentially important PBRM1-dependent change in association with H3K9me3 (Fig. 4e,f). Given the importance of this mark in pericentromeric heterochromatin, and the changes in H3K9me3 staining detected in PBRM1 deficient cells (Fig. 2), we explored this relationship further. First, we find that the number of KO specific *k*- mers that intersect with H3K9ac enriched *k*-mers is lower than for the PBRM1 specific subset, similar to what is seen with H3K4me3, and instead, there is a greater number of KO specific *k*- mers that intersect with H3K9me3 enriched *k*-mers. (Fig. 4g and Supplementary Fig. 9a).

To look at this more quantitatively, we analysed the proximity of these by plotting the relative distance between the mapping locations of subsets of SWI/SNF *k*-mers and histone PTM *k*- mers as a cumulative fraction^24^. To test the utility of this approach, we first analysed the promoter-associated H3K4me3 mark and find that, as expected, more PBRM1 specific mapping locations are closer to those of H3K4me3 than those of the PBRM1 non-specific *k*- mers. (Fig. 4h and Supplementary Fig. 9b).

Interestingly, KO specific *k*-mer mapping locations tend to be further away from those of H3K4me3 *k*-mers (Fig. 4h and Supplementary Fig. 9b). In contrast, these are much closer to mapping locations that are normally enriched in H3K9me3 and further away from those enriched in H3K9ac (Fig. 4h). These data suggest that in the absence of PBRM1, SMARCA4 inappropriately localizes to what are normally heterochromatic regions flanking centromeres.

### PBRM1 KO cells are sensitive to mitotic perturbation

We next wanted to test whether the changes in structure at centromeres and pericentromeric regions where PBRM1 is bound lead to sensitivity to mitotic perturbation when PBRM1 is absent. We tested this and found PBRM1 KO cells are sensitive to chronic CDK1 inhibition compared with parental RPE1 cells (Fig. 5a,b). We also treated cells with an acute dose of CDK1 inhibitor (Fig. 5c) and monitored the presence of aberrant mitotic events after release.

**Figure 5.**
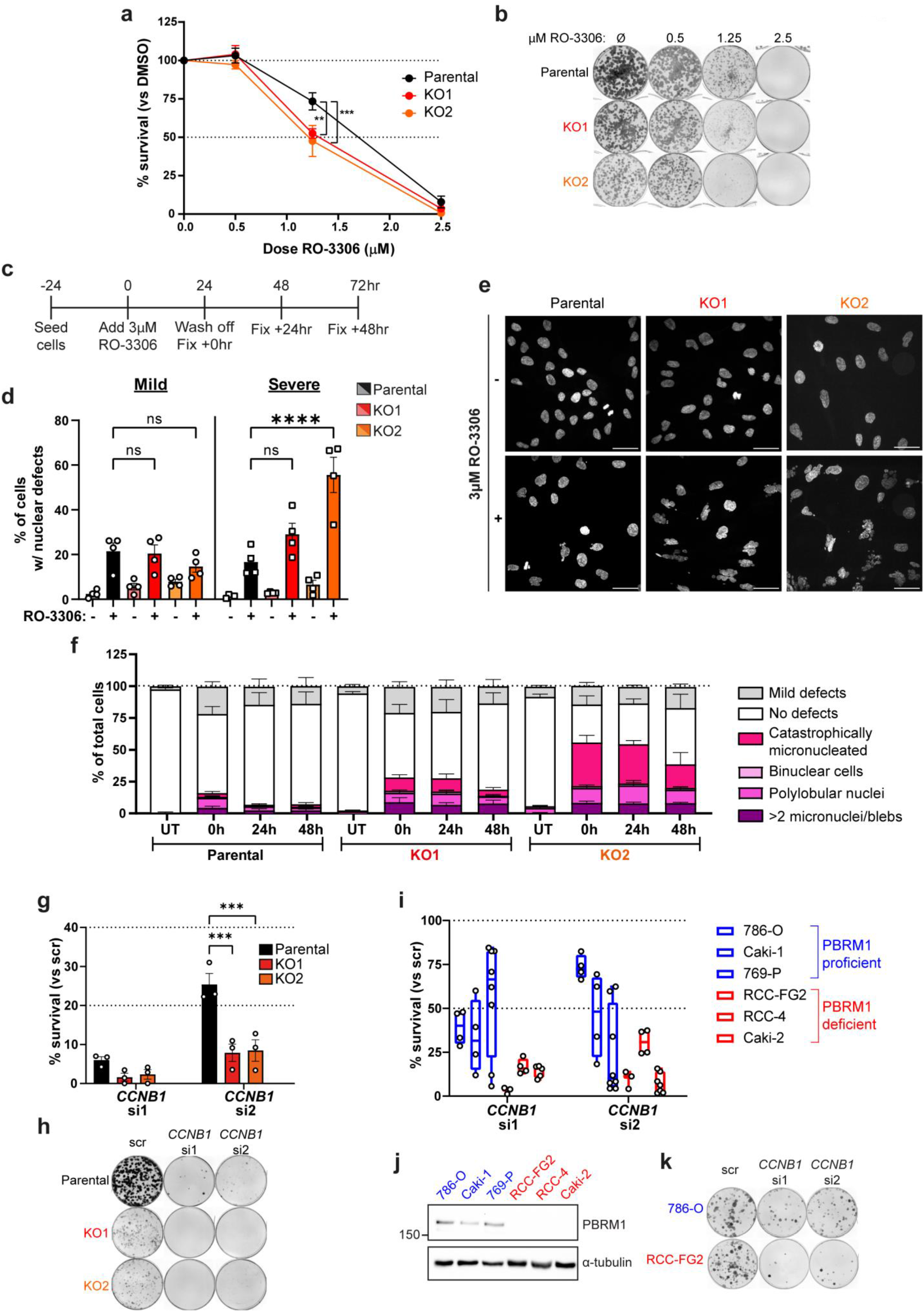
PBRM1 KO cells are sensitive to mitotic perturbation. **(a)** Clonogenic survival of RPE1 parental and PBRM1 knockouts when treated with increasing doses of the CDK1 inhibitor RO-3306 relative to DMSO-only treated cells. *n*=4, mean±SEM, and data were analysed using a 2way ANOVA with Dunnett’s test, \*\**p*<0.01, \*\*\**p*<0.001. **(b)** Representative image of colony formation from clonogenic survival assay in (a). **(c)** Experimental outline for quantifying nuclear defects induced by CDK1 inhibition. RPE1 parental or PBRM1 knockout cells were treated with either 3µM RO-3306 or DMSO alone for 24 hours, followed by wash-off of the drug and fixation at 0, 24, and 48hours after wash-off for immunofluorescence imaging. Nuclear defects were only quantified in interphase cells. **(d)** Quantification of the % of total cells imaged with either mild (left) or severe (right) nuclear defects after 24 hours treatment with either DMSO alone (-) or 3µM RO-3306 (+). *n*=4, mean±SEM, and data were analysed by 2way ANOVA with Dunnett’s test, \*\*\*\**p*<0.0001. 600-1500 nuclei were analysed for each condition. **(e)** Representative images from data in (D), showing changes in nuclear morphology after 24 hours treatment with DMSO alone (-) or 3µM RO-3306 (+). Scale bar corresponds to 40µm. **(f)** Details of nuclear morphology of parental or PBRM1 knockout cells treated with DMSO alone (UT) or 0, 24, and 48 hours after treatment with 3µM RO-3306 for 24 hours. Nuclei were defined as having no defects, mild defects (1-2 micronuclei or nuclear blebs per nucleus), or developing one of the severe nuclear defects, i.e. having >2 micronuclei/nuclear blebs per nucleus, or exhibiting a catastrophically micronucleated, binuclear, multinucleated, or polylobular phenotype. **(g)** Clonogenic survival of RPE1 parental and PBRM1 knockouts after *CCNB1* RNAi with two independent *CCNB1* siRNAs, normalised to survival after treatment with a scramble siRNA. Points correspond to independent biological replicates, *n*=3, mean±SEM, and data were analysed by 2way ANOVA with Dunnett’s test, \*\*\**p*<0.001. **(h)** Representative image of colony formation from clonogenic survival assay in (g). **(i)** Clonogenic survival of a panel of renal cell carcinoma (RCC) cell lines, 3 of which are PBRM1 proficient (blue) and 3 of which are PBRM1 deficient due to loss-of-function mutations (red). Clonogenic survival was measured after *CCNB1* RNAi with two independent *CCNB1* siRNAs, normalised to survival after treatment with a scramble siRNA. Points correspond to independent biological replicates, *n*=3-8 (depending on cell line), boxplot shows min to max with line at median. **(j)** Western blotting for PBRM1 in PBRM1 proficient and deficient RCC cell lines. α-tubulin is used as a loading control. **(k)** Representative images showing colony formation in one PBRM1 proficient (786-O) and one PBRM1 deficient (RCC-FG2) cell line from the clonogenic survival assay in (i).

PBRM1 KO cells show a substantial increase in the number of nuclear abnormalities (Fig. 5d-f, Supplementary Fig. 10a,b).

We also found PBRM1 KO cells are selectively sensitive to depletion of CCNB1 (Fig. 5g,h and Supplementary Fig. 10c). 786-O derived PBRM1 KO clones, however, which do not show downregulation of centromere- or pericentromere-associated proteins (Supplementary Fig. 2d, and Supplementary Fig.12a,b), also do not exhibit sensitivity to CCNB1 depletion (Supplementary Fig. 12c,d). We further tested this using a panel of six renal cancer cell lines, in which three have loss of function mutations in PBRM1, and we found that PBRM1 deficient cells are considerably more sensitive to CCNB1 depletion than the PBRM1 proficient cell lines (Fig. 5i-k, and Supplementary Fig.S10d,e), suggesting potential clinical implications. These data suggest that loss of PBRM1 reduces the ability to cope with mitotic perturbations.

### PBRM1 KO cells display sensitivity to Mps1 inhibition *in vitro* and *in vivo*

Cells with aberrant centromeres rely on the activity of the spindle assembly checkpoint (SAC) if these changes impact on kinetochore-microtubule attachments^25^. We therefore tested whether cells with PBRM1 loss are sensitive to inhibition of the Mps1 kinase that regulates the SAC^25^. Using three different Mps1 inhibitors, we found that RPE1-derived PBRM1 KO cells are modestly but significantly sensitive when compared with the parental cells (Fig. 6a-c, and Supplementary Fig. 11a-c). In contrast, we found no substantial sensitivity to Mps1 inhibitors in the 786-O derived PBRM1 KO cells (Supplementary Fig. 12e-h), supporting the link between downregulation of centromere-associated proteins and perturbation of mitosis.

**Figure 6.**
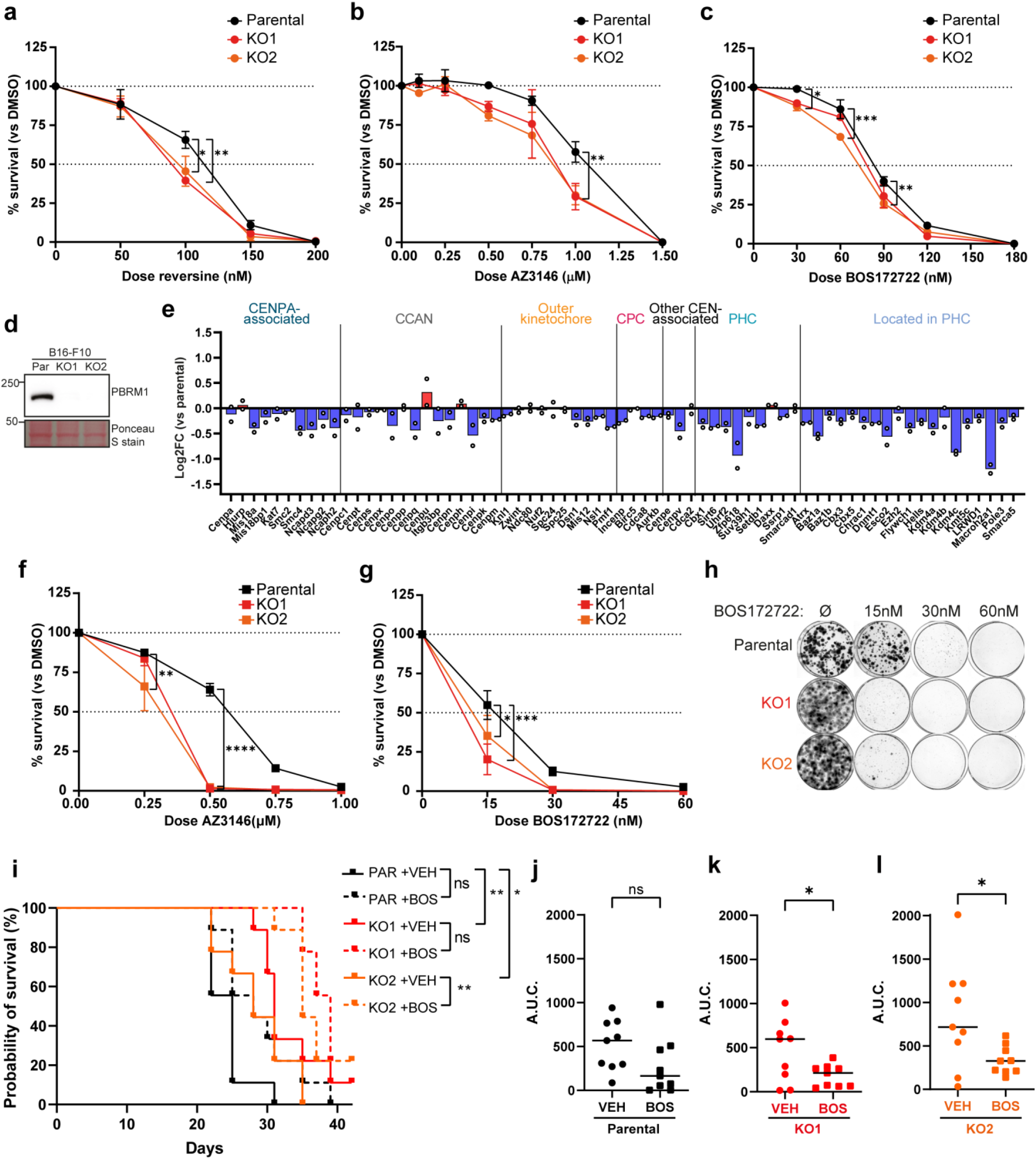
PBRM1 KO cells display sensitivity to Mps1 inhibition in vitro and in vivo. **(a)-(c)** Clonogenic survival of RPE1 parental and PBRM1 knockout cells after treatment with increasing doses of the Mps1 inhibitors (a) reversine, (b) AZ3146, and (c) BOS172722. *n*=3-4, mean±SEM, and data were analysed by 2way ANOVA with Dunnett’s test, \**p*<0.05, \*\**p*<0.01, \*\*\**p*<0.001. **(d)** Western blotting for PBRM1 in B16-F10 parental and PBRM1 knockout cells. Ponceau S stain is used to show whole protein levels. **(e)** Median Log2FC of annotated peri/centromere proteins in B16-F10 PBRM1 knockouts compared to parental cells. Points correspond to individual knockout clones from one of two independent biological replicates. **(f)-(g)** Clonogenic survival of B16-F10 parental and PBRM1 knockout cells after treatment with increasing doses of Mps1 inhibitors (f) AZ3146 and (g) BOS172722. *n*=4, mean±SEM, and data were analysed by 2way ANOVA with Dunnett’s test, \**p*<0.05, \*\**p*<0.01, \*\*\**p*<0.001, \*\*\*\**p*<0.0001. **(h)** Representative image of colony formation from clonogenic survival assay in (g). **(i)** Kaplan-Meier survival curve showing survival of mice with tumours derived from B16-F10 parental or PBRM1 knockout cells, treated with either vehicle (VEH) or BOS172722 (BOS). *n*=9 mice per condition, and survival was compared using the logrank (Mantel-Cox) test, \**p*<0.05, \*\**p*<0.01. **(j)-(l)** Area under the curve (A.U.C.) was calculated based on tumour growth up to the latest timepoint where all mice were still surviving in the indicated cell line – Day 21 for tumours derived from (j) B16-F10 parental and (l) PBRM1 KO2 cells, and Day 27 for (k) PBRM1 KO1 cells. Significance was determined using an unpaired t-test, \**p*<0.05.

To test whether this has clinical potential, we created two PBRM1 KO clones in the mouse melanoma B16-F10 cell line for *in vivo* studies (Fig. 6d and Supplementary Fig. 11d). We first analysed the proteome of these cells by mass spectrometry, and strikingly, we found downregulation of centromere- and pericentromere-associated proteins (Fig. 6e), suggesting that this is a conserved feature of PBRM1 loss in at least one other species. We next tested the response to Mps1 inhibitors and found the survival of the mouse PBRM1 KO cell lines was significantly lower than the parental B16-F10 cells (Fig. 6f-h, and Supplementary Fig. 11e), consistent with a functional impact of altered centromere associated proteins.

We used these cell lines to perform *in vivo* studies with Mps1 inhibitors. Mice were dosed twice weekly with the BOS172722 Mps1 inhibitor over the course of 42 days, at which point the study was terminated (Supplementary Fig. 11f). Mps1 inhibitor treatment had a marginal and statistically insignificant impact on survival of mice bearing B16-F10 parental cell line derived tumours (Fig. 6i and Supplementary Fig. 11g). In contrast, survival was improved following Mps1 inhibitor treatment in mice bearing the PBRM1 KO derived tumours (Fig. 6i, and Supplementary Fig. 11h,i). Moreover, tumour growth of PBRM1 KO-derived tumours, but not that of parental B16-F10 derived tumours, was significantly reduced following treatment with Mps1 inhibitors (Fig. 6j-l and Supplementary Fig. 11j). Together, these data suggest that reliance on the SAC in PBRM1 KO cancers represents a therapeutic vulnerability that can be clinically exploited.

### Loss of PBRM1 leads to centromere fragility

These results raised the possibility that cells lacking PBRM1 display centromere fragility. To test this, we examined the patterns of sister chromatid exchanges (SCEs) in RPE1 parental and PBRM1 KO cell lines in the absence of any perturbations. We found a slightly elevated number of total SCEs in the PBRM1 KO clones compared with the parental RPE1 cells (Fig. 7a,c and Supplementary Fig. 13a). Notably, when we scored the location of the exchanges, we found that they were more likely to localise to centromeres in the PBRM1 KO cells (Fig. 7b,c, and Supplementary Fig. 13b), consistent with an increased vulnerability at centromeres.

**Fig. 7.**
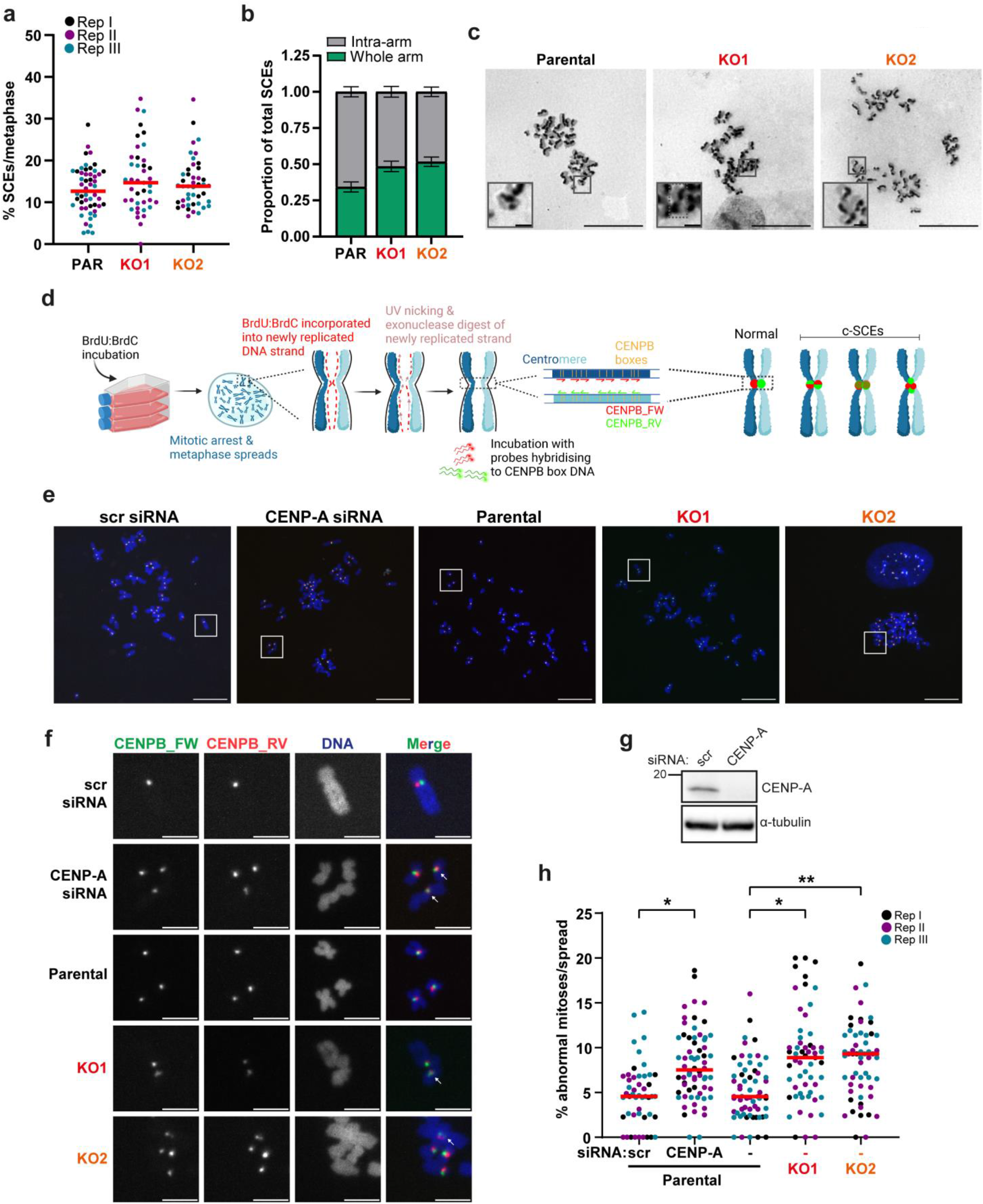
Loss of PBRM1 leads to centromere fragility. **(a)** Quantification of the % of chromosomes in metaphases with sister chromatid exchanges (SCEs) in RPE1 parental or PBRM1 knockout cells. *n*=3, with different colours corresponding to independent biological replicates and points indicating individual metaphases, line at median, and data were analysed using 1way ANOVA with Dunnett’s test on the median of each biological replicate. **(b)** Quantification of the proportion of total SCEs which were intra-arm (grey) versus whole arm (green) exchanges per metaphase, in RPE1 parental and PBRM1 knockout cells. Data are presented as mean±SEM, *n*=3. **(c)** Representative images of metaphases in RPE1 parental and PBRM1 knockout cells stained with Giemsa to visualise SCEs. Scale bar corresponds to 20µm. Zoomed images showed intra-arm (RPE1 parental) or whole arm (KO1 & KO2) SCEs, and scale bar corresponds to 5µm. Dotted line zoomed KO1 image highlights a single chromosome with whole arm exchange. **(d)** Schematic showing Cen-CO-FISH workflow, and methods of quantifying normal versus aberrant centromeres. **(e)** Representative metaphase spreads from Cen-CO-FISH experiments, with DNA (blue) stained with DAPI, and centromeres hybridised with FISH probes against CENP-B. Scale bars correspond to 20µm. White boxes correspond to zoomed images in (f). **(f)** Zoomed images of representative individual centromeres from Cen-CO-FISH experiments. White arrows indicate aberrant centromeres. Scale bars correspond to 5µm. **(g)** Western blotting for CENP-A after treatment of RPE1 parental cells with either scramble or CENP-A siRNA for subsequent Cen-CO-FISH analyses. α-tubulin was used as loading control. **(h)** Quantification of sister chromatid exchanges at centromeres, defined as the percentage of chromosomes with aberrant centromeres in each metaphase spread (% abnormal mitoses). At least 49 metaphases were quantified for each condition. *n*=3, with different colours corresponding to independent biological replicates, line at median, and data were analysed using 2way ANOVA with Dunnett’s test, \**p*<0.05, \*\**p*<0.01.

To look more directly at centromere fragility, we made use of an assay in which centromeres are labelled in a strand specific manner to identify sister chromatid exchanges and other centromere-specific rearrangements (termed Cen-CO-FISH^11^; Fig. 7d). As a positive control, we depleted CENPA, which protects centromere integrity^11^. Strikingly, we found that the PBRM1 KO clones showed significantly elevated levels of aberrant centromere signals when compared with the parental RPE1 cells, at a level similar to that seen in CENPA-depleted cells (Fig. 7e-h). Our mapping data suggests that other SMARCA4-containing complexes are bound at flanking pericentromeric regions (Fig. 3). We were therefore interested to see whether other SWI/SNF complexes are involved in protecting genome stability at centromeres. We performed the Cen-CO-FISH assay in RPE1 cells with CRISPR-mediated KO of the ARID1A subunit of BAF (Supplementary Fig. 13c), which is also commonly mutated in cancer^8^. Notably, in contrast to the effect of PBRM1 loss, we found no detectable difference in aberrant centromere signals in the ARID1A KO cells when compared with parental RPE1 (Supplementary Fig. 13d,e). We additionally performed the assay in a second cell line background and again find elevated levels of aberrant centromere signals in the PBRM1 KO clones when compared with isogenic controls (Supplementary Fig. 13f,g). Collectively, these data indicate that loss of PBRM1 leads to substantially increased genome instability at centromeres, even in the absence of any perturbations.

## Discussion

Here, we find that changes to the structure of centromeres and centromere-associated protein levels are conserved features of PBRM1-deficient cells. We find PBRM1 directs SMARCA4 to specific locations in chromatin flanking centromeres. In the absence of PBRM1, cells show mitotic defects and vulnerability to inhibition of the spindle assembly checkpoint, which has the potential to be therapeutically exploited. Most notably, we find centromere fragility in PBRM1-deficient cells, even in the absence of any perturbation.

Together, these data are consistent with a model (Supplementary Fig. 13h) in which PBRM1- containing PBAF complexes work in the transition arms flanking centromeres to help to create a chromatin substrate that promotes assembly of protein complexes and structures. In the absence of PBRM1, the integrity of these structures is compromised, leading to mitotic vulnerability, dependence on the spindle assembly checkpoint, and inappropriate recombination between repetitive sequences at the centromere.

When mapping SMARCA4 to pericentromeric regions, we find that a subset of binding sites are lost when PBRM1 is deficient, suggesting that PBRM1 is important for directing SMARCA4-containing complexes to these sites. This raises the possibility that there are acetylated substrates at these sites acting as beacons for the PBRM1 bromodomains.

Perhaps more interestingly, we find that SMARCA4 localizes to new sites when PBRM1 is deficient, and many of these sites are normally enriched in H3K9me3. We also find that H3K9me3 patterns are altered when PBRM1 is deficient, and that PBRM1 localization anti-correlates with H3K9me3 enriched regions. These data are consistent with the idea that PBRM1-containing PBAF complexes help to establish appropriate patterns of H3K9me3, which is important for the spatial organization of chromatin^26^. It was recently reported that DSBs exist in centromeres^6^. We previously showed that PBRM1 influences cohesin dynamics, both at centromeres^10^ and DSBs, to help promote repair^27^. H3K9me3 can also influence centromere function and DSB repair^3, 28^, and there is an interplay between H3K9me3 and cohesin dynamics^29, 30^. Therefore, one attractive possibility is that PBRM1 could promote stability at centromeres and pericentromeric regions by modulating the chromatin environment (via H3K9me3 patterns and cohesin dynamics) to prevent inappropriate DSB processing, which would lead to aberrant recombination events between repeats.

Importantly, PBRM1 loss was recently identified as a genetic determinant for a chromosome instability signature that is associated with whole arm or whole chromosome changes^31^. This is consistent with the role we identify here in preventing centromere fragility. Notably, ARID1A, which is frequently mutated in cancer, was not identified as a determinant for chromosomal instability, and consistently, we find no impact of ARID1A on centromere fragility in our assays. This suggests that, while BAF may also be bound in regions in or around centromeres, it doesn’t provide the same function as PBAF complexes to prevent centromere fragility. Importantly, we observe PBRM1-dependent centromere fragility in the absence of any exogenous perturbation, suggesting that this fundamental feature of PBRM1 loss is a critical activity that contributes to tumourigenesis and cancer progression.

## Methods

### Cell culture

All cell lines were obtained from ATCC unless otherwise specified. hTERT-RPE1 cells were cultured in Dulbecco Modified Minimal Essential Medium (DMEM)/F-12 (Sigma) supplemented with 10% FBS (Gibco), 200µM glutamax (Gibco), 0.26% sodium bicarbonate (Gibco), and 1% penicillin/streptomycin (P/S)(Sigma). 1BR3-hTERT (Gift from Penny Jeggo, Sussex University), Hek293TN, U2OS, A375, RCC4-VO (European Collection of Authenticated Cell Cultures, ECACC), and B16-F10 cells were cultured in DMEM supplemented with 10% FBS and 1% P/S. MOC-2 cells were cultured in DMEM supplemented with 5% FBS, 100µM glutamax, and 1% P/S. 786-O, 769-P, A704, and RCC- FG2 (Cell Lines Service, CLS GmbH) cells were cultured in RPMI 1640 medium (Sigma) supplemented with 10% FBS and 1% P/S. Caki-1 and Caki-2 cells were cultured in McCoy’s 5A modified medium (Gibco) supplemented with 10% FBS and 1% P/S. All cells were maintained at 37°C in a humified incubator with 5% CO2 and were regularly tested for mycoplasma contamination.

### Drug treatments

For clonogenic survival and SRB assays, cells were treated with a range of concentrations of RO-3306 (Merck), reversine (Sigma), AZ3146 (Sigma), or BOS172722 (MedChemExpress). For immunofluorescence experiments, cells were treated with 3µM RO-3306. At this dose, partial inhibition of CDK1 is seen, perturbing mitotic timing and progression, but not causing the G2 arrest seen at higher doses of RO-3306^32^. For in vivo experiments, BOS172722 ditosylate salt was dissolved in vehicle (10% DMSO and 1% Tween-20 in 89% saline).

### CRISPR/Cas9-mediated gene knockout

For PBRM1 knockout in a range of cell lines and ARID1A knockout in RPE1, sgRNAs (Sigma) targeting PBRM1 or ARID1A Exon 1 were cloned into the pSpCas9(BB)-2A-GFP (PX458) construct. For SMARCA4 knockout in RPE1 cells, two sgRNAs targeting Exon 2 of SMARCA4 were cloned into the CRISPR-Cas9 nickase pSpCas9n(BB)-2A-GFP (PX461) construct. gRNAs are listed in Supplementary Table S1. PX458 and PX461 plasmids were a gift from Professor Feng Zhang (Addgene plasmids #48138 and #48140 respectively).

Plasmids were transfected into cells using Lipofectamine 3000 (Invitrogen), according to the manufacturer’s instructions. 48 hours after transfection, GFP- (PX458) or mRuby-positive (PX461) cells were single cell sorted into 96-well plates using a FACSAria™ III sorter (BD).

Resulting clones were screened for loss of PBRM1 or SMARCA4 using western blotting, followed by Sanger sequencing (Eurofins) of the targeted genomic region, as well as immunofluorescence microscopy and proteomic profiling. The PBRM1 knockouts in RPE1 and 1BR3 were described in^9^.

### siRNA mediated gene depletion/RNA interference

100µM of the specified siRNAs (Dharmacon) were reverse transfected into cells using Lipofectamine RNAiMAX (Invitrogen), according to the manufacturer’s instructions. Multiple single siRNAs or pools containing a mix of 4 independent siRNAs were used as indicated in Fig. legends. Corresponding single or pooled non-targeting siRNAs (Dharmacon) were used as control. Cells were assayed 24-72 hours after transfection.

### Clonogenic survival assay

Cells in culture were diluted to the appropriate density (300-1,000 cells per dish, depending on the cell line) and seeded onto 6cm dishes in triplicate. For survival after drug treatments, drugs were added 24 hours after seeding. For clonogenic survival after RNAi, cells were reverse transfected as described with the appropriate siRNA 24 hours before seeding. Cells were incubated in culture for 10-21 days (depending on cell line), after which media was aspirated and colonies were fixed and stained with 1% methylene blue in 70% methanol for 1 hour at room temperature with gentle rocking. Dishes were washed extensively with water and allowed to dry overnight. Colony number was counted using a Digital Colony Counter (Stuart), and survival was defined as the % of colonies in treated conditions versus control conditions (vehicle only or control siRNA for drug treatments and RNAi respectively).

### Proliferation assays

For cell growth assays, 1x10^5^ cells were seeded to 6 well plates. Every 24 hours, triplicate wells were trypsinised and counted, up to a total of 96 hours, to quantify the speed of proliferation. For hTERT-RPE1 cells, cells were seeded in triplicate into 96-well plates, and the rate of proliferation was quantified on the Incucyte SX5 (Sartorius) using phase contrast images of cells taken every 4 hours, up to a total of 116 hours.

### Flow cytometry

For flow cytometric analysis of cell cycle distribution, growth medium was collected, and cells were trypsinised and combined with cells suspended in growth medium. Cells were pelleted by centrifugation and washed once with PBS. After removal of most supernatant, cells were gently resuspended in remaining PBS and fixed by addition of 70% ethanol dropwise with gentle vortexing and incubated at −20°C overnight. Cell suspensions were centrifuged for 5 minutes at 300 x g and the ethanol removed. Pellets were washed twice in cold PBS and then resuspended in the appropriate volume of PBS containing 5µg/mL propidium iodide (Sigma) and 0.1mg/mL RNase A. Cells were incubated at 37°C for 30 minutes, protected from light. DNA content of at least 10,000 cells per condition was detected on a BD LSR II or BD FACSymphony A5 (BD Biosciences). Cell debris and doublets were removed by gating and cell cycle phases were quantified using FlowJo software (v10.8.1).

### Immunofluorescence microscopy

Cells for immunofluorescence (IF) imaging were seeded onto glass coverslips in 6 well plates and assayed. Cells were fixed with 100% ice-cold methanol for 15 minutes at −20°C, followed by rehydration with four 5-minute washes in PBS. Samples were blocked with 1% BSA in PBS for 1 hour at room temperature and incubated overnight at 4°C in primary antibody diluted in blocking solution. Following washes in PBS, cells were incubated with the appropriate secondary antibody and DNA stain Hoechst 33342 (Sigma) for 2 hours at room temperature. Cells were then washed in PBS, mounted onto frosted glass microscope slides with ProLong Gold (Thermo Fisher Scientific), and cured overnight. Antibodies used for immunofluorescence imaging are indicated in Table S2. Cells were imaged with an Advanced Spinning Disk or Super Resolution SoRa Spinning Disk Confocal microscope with SlideBook imaging software (3i). 1.02µm Z-stacks were imaged using a 40x or 63x oil objective, or 0.27µm Z-stacks when using the SoRa spinning disk, and exported for analysis as maximum intensity projections.

### Fluorescence in situ hybridisation (FISH)

Cells were trypsinised and washed once in PBS. Washed cells were then fixed by adding ice-cold fixative solution (3:1 methanol:acetic acid) dropwise with gentle vortexing. Cells were incubated for 15 minutes at room temperature, centrifuged, and subjected to a second round of fixation. Cells were then resuspended in 30µL fixative solution, dropped onto a glass microscope slide, and allowed to dry overnight. After drying, cells were rehydrated in 2x sodium chloride and sodium citrate (SSC) buffer for 2 minutes followed by dehydration in an ethanol series (70, 80, and 95%) for 2 minutes each. The chromosome 2 and chromosome 10 α-satellite FISH probes (Cytocell) were denatured and hybridised with slides according to the manufacturer’s protocol. After hybridisation, cells were washed in 0.25x SSC for 2 minutes at 72°C and 2x SSC with 0.05% Tween-20 for 30 seconds at room temperature. Slides were drained and mounted with Vectashield Antifade mounting medium with DAPI (Vector labs).

### Metaphase spreads

Cells in culture were treated with 0.1µg/mL colcemid (Gibco) for 3-5 hours to accumulate cells in mitosis. Cells were trypsinised, saving floating cells and PBS washes in a falcon tube, and the supernatant and resuspended cells were pelleted, washed with PBS, and slowly resuspended in 10mL of pre-warmed hypotonic solution (75mM potassium chloride) and incubated for 30 minutes at 37°C with intermittent gentle inversion. 200µL fresh ice-cold fixative solution (3:1 methanol:acetic acid) was added to solution directly before centrifugation at 175 x g for 5 minutes. The majority of the supernatant was removed, leaving less than 0.3mL, and the pellet was gently resuspended in the remaining supernatant by tapping. Cells were fixed by adding 10mL fixative solution dropwise with gentle vortexing and incubated at room temperature for 15 minutes. Cells were washed once with fixative solution and resuspended in the appropriate amount of fixative solution to result in a cell concentration of approximately 1x10^6^ cells/mL. Drops of approximately 20µL were dropped onto humid glass slides from a height and air-dried overnight.

### Sister chromatid exchange (SCE) assay

SCE assay was performed as described previously^33, 34^. Briefly, cells were labelled for two cell cycles, calculated based on doubling time, with 20µM BrdU. Metaphases were fixed and dropped onto slides, as described above. After drying, slides were immersed in 10µg/mL Hoechst-33258 (Sigma) for 20 minutes at room temperature. After rinsing in ddH2O, slides were exposed to 365nm UV light for 30 minutes, immersed in 2x SSC buffer. Slides were then immersed in preheated 2x SSC at 50°C for 1hr, before being incubated in 10% Karyo- MAX^TM^ Giemsa stain solution in Sorenson buffer (pH=6.8) for 30 minutes at room temperature. After incubation, slides were rinsed with ddH2O and allowed to dry, followed by mounting with DPX medium (Sigma). Metaphases were imaged on an RGB colour camera using 3i imaging software.

### Cen-CO-FISH

Cen-CO-FISH was performed as described^35^. Briefly, cells were labelled for a single cell cycle, calculated based on doubling time, with 7.5mM 5’-Bromodeoxyuridine (BrdU)(Sigma) and 2.5mM 5’-Bromodeoxycytidine (BrdC)(MP Biomedicals). Cells were trypsinised and metaphase spreads were prepared as described above, ensuring that metaphase spreads were protected from light when drying. The following day, slides were rehydrated in PBS, and treated with 0.5mg/mL RNase A (Sigma) for 10 minutes at 37°C. Slides were then stained with 1µg/mL Hoechst-33258 (Sigma) in 2x SSC buffer. Slides covered in 2x SSC buffer were exposed to 365nm UV irradiation for 30 minutes using a UV lamp (Analytik Jena). Following UV nicking, DNA was digested using Exonuclease III (Promega) dissolved in the appropriate buffer according to the manufacturer’s instructions, twice for 10 minutes each. Slides were washed, dehydrated using an ethanol series (70, 90, and 100%), and dried overnight. The following day, slides were hybridised with peptide nucleic acid (PNA) probes against the CENP-B box (PNABio) diluted to a 50nM concentration in hybridisation solution (10mM Tris-HCl (pH=7.4) and 0.5% blocking reagent (Roche) in 70% formamide) and heated to 60°C for 10 minutes directly before use. Slides were first hybridised in a dark hybridisation chamber for 2 hours at room temperature with the forward CENP-B probe (PNA Bio, #3004), washed for 30 seconds in Wash Buffer #1 (10mM Tris-HCl (pH=7.4) and 0.1% BSA in 70% formamide), and incubated for 2 hours with the reverse CENPB probe (PNA Bio, #F3009). Slides were then washed twice in Wash Buffer #1 for 15 minutes each with gentle rocking, followed by three washes in Wash Buffer #2 (0.1M Tris-HCl (pH=7.4), 0.15M NaCl, and 0.1% Tween-20), including 1µg/mL DAPI (Sigma) in the second wash.

Slides were dehydrated in an ethanol series (70, 90, and 100%), air dried, and mounted using ProLong Gold. Slides were imaged as before on an Advanced Spinning Disk confocal microscope but instead imaging 0.2µm Z-stacks and were exported as maximum intensity projections.

### Whole protein extraction

Cell pellets were resuspended in the appropriate volume of lysis buffer (10% glycerol, 50mM Tris-HCl (pH=7.4), 0.5% NP-40, 150mM NaCl) containing 0.25U/µL benzonase nuclease (Sigma), 1x cOmplete™ EDTA-free protease inhibitor cocktail (Roche), and 1x PhosSTOP™ phosphatase inhibitor cocktail (Roche). Cells were lysed on ice for 45 minutes followed by centrifugation for 15 minutes at 16,000 x g at 4°C. The resulting supernatant containing whole cell protein extracts was collected. For the SMARCA4 western blot in Fig. S9, whole cell extracts were obtained using urea buffer as described previously ^36^. Protein concentration was estimated using a Bradford assay (Bio-Rad) according to the manufacturer’s protocol.

### Western blotting

For each western blot, approximately 30µg of protein extract was combined with LSB containing 5% β-mercaptoethanol and boiled for 10 minutes at 95°C to denature proteins. Proteins were separated via SDS-polyacrylamide gel electrophoresis and transferred to 0.2µm nitrocellulose membranes (Fisher Scientific). Successful protein transfer was confirmed by incubating membranes in Ponceau S solution (Sigma) for 5 minutes with rocking and membranes were blocked in 5% milk in TBS buffer containing 0.1% Tween-20 for 1 hour with gentle rocking. Membranes were incubated in the appropriate antibody diluted in blocking buffer at 4°C overnight, washed 3 times with TBS buffer containing 0.1% Tween-20, and incubated in the appropriate horseradish peroxidase (HRP)-conjugated secondary antibody diluted in blocking buffer for 1 hour at room temperature (except for β-actin antibody, which is already HRP-conjugated). Proteins were visualised on an iBright CL750 imager (Thermo Fisher Scientific) using Immobilon Forte HRP substrate for chemiluminescence. Details of the primary and secondary antibodies used for western blotting in this study are detailed in Supplementary Table S2.

### RT-qPCR

RNA was extracted from cells using an RNeasy Mini Kit (Qiagen) according to the manufacturer’s protocol. 0.5µg of RNA was then reverse transcribed to cDNA with the High-Capacity cDNA Reverse Transcription kit (Applied Biosystems) according to the manufacturer’s protocol. 4ng of cDNA was used for each qPCR reaction, along with the indicated forward and reverse primers (Supplementary Table S1) at 200nM concentration, and Power SYBR green PCR master mix (Applied Biosystems). Samples were run in triplicate in MicroAmp Fast Optical 96-well plates (Applied Biosystems) on a StepOne Plus Real-Time PCR system (Applied Biosystems), according to the manufacturer’s protocol. cDNA levels were compared using the Ct (comparative cycle) method, and GAPDH and PPIA were used as housekeeping genes to normalise data.

### Proteomics analysis – sample preparation

Cell pellets were lysed in 150μL buffer containing 1% sodium deoxycholate (SDC), 100 mM triethylammonium bicarbonate (TEAB), 10% isopropanol, 50mM NaCl and Halt protease and phosphatase inhibitor cocktail (100x) (ThermoFisher Scientific) on ice, assisted with probe sonication, followed by heating at 90°C for 5 min and re-sonication for 5 sec. Protein concentration was measured with the Quick Start Bradford protein assay (Bio-Rad) according to manufacturer’s instructions. Protein aliquots of 60μg or 100μg (for TMTpro and TMT11plex respectively) were reduced with 5 mM tris-2-carboxyethyl phosphine (TCEP) for 1 hour at 60°C and alkylated with 10 mM iodoacetamide (IAA) for 30 min in the dark, followed by overnight digestion with trypsin at a final concentration of 75ng/μL (Pierce).

Peptides were labelled with the TMT-11plex or TMTpro reagents (ThermoFisher Scientific) according to manufacturer’s instructions. The TMT labelled peptide pool was acidified at 1% formic acid, the precipitated SDC was removed by centrifugation, and the supernatant was SpeedVac dried. Peptides were fractionated with high-pH Reversed-Phase (RP) chromatography with the XBridge C18 column (2.1 x 150mm, 3.5 μm) (Waters) on a Dionex UltiMate 3000 HPLC system. Mobile phase A was 0.1% (v/v) ammonium hydroxide and mobile phase B was acetonitrile, 0.1% (v/v) ammonium hydroxide. Peptides were fractionated at a flow rate of 0.2 mL/min using the following gradient: 5 min at 5% B, for 35 min gradient to 35% B, gradient to 80% B in 5 min, isocratic for 5 minutes and re-equilibration to 5% B. Fractions were collected every 42 sec, combined in 28 fractions and SpeedVac dried.

### Proteomics analysis – LC-MS and protein quantification

LC-MS analysis was performed on a Dionex UltiMate 3000 UHPLC system coupled with the Orbitrap Lumos Mass Spectrometer (ThermoFisher Scientific). Peptides were loaded onto the Acclaim PepMap 100, 100μm × 2cm C18, 5μm, trapping column at 10μL/min flow rate. Peptides were analysed with the EASY-Spray C18 capillary column (75μm × 50cm, 2μm). Mobile phase A was 0.1% formic acid and mobile phase B was 80% acetonitrile, 0.1% formic acid. Peptides were separated over a 90 min gradient 5%-38% B at a flow rate of 300 nL/min. Survey scans were acquired in the range of 375-1,500 m/z with a mass resolution of 120K. Precursors were selected in the top speed mode in cycles of 3 sec and isolated for CID fragmentation with quadrupole isolation width 0.7 Th. Collision energy was 35% with AGC 1×104 and max IT 50 ms. Quantification was obtained at the MS3 level with HCD fragmentation of the top 5 most abundant CID fragments isolated with Synchronous Precursor Selection (SPS). Quadrupole isolation width was 0.7 Th and collision energy was 65%. The HCD MS3 spectra were acquired for the mass range 100-500 with 50K resolution. Targeted precursors were dynamically excluded for further fragmentation for 45 seconds with 7 ppm mass tolerance. The mass spectra were analysed in Proteome Discoverer 2.2 or 2.4 (ThermoFisher Scientific) with the SequestHT search engine for protein identification and quantification. The precursor and fragment ion mass tolerances were set at 20 ppm and 0.5 Da respectively. Spectra were searched for fully tryptic peptides with maximum 2 missed cleavages. TMT6plex or TMTpro at N-terminus/K and Carbamidomethyl at C were selected as static modifications. Oxidation of M and Deamidation of N/Q were selected as dynamic modifications. Peptide confidence was estimated with the Percolator node and peptides were filtered at q-value<0.01 based on target-decoy database search. All spectra were searched against reviewed UniProt Homo sapiens protein entries. The reporter ion quantifier node included a TMT11plex or TMTpro quantification method with an integration window tolerance of 15 ppm at the MS3 level. Only unique peptides were used for quantification, considering protein groups for peptide uniqueness. Only peptides with average reporter signal-to-noise>3 were used for protein quantification.

### Statistical analysis of proteomics data

Log2 ratios against the parental (PBRM1 WT) cells were computed for each PBRM1 knockout (KO) clone, per cell line, using the sum-normalised abundances exported from Proteome Discoverer. For visualising effects on each cell line (Fig. 1G-I and S2C-D), the median log2(KO/WT) was calculated from replicate clones or biological repeats and was further normalised by subtracting column median. Proteins detected in least one parental and PBRM1 knockout cell line were used for GSEA analysis (v4.3.2)^37^. The log2 ratio of classes was used as the metric for ranking genes, and normalised abundances from proteomic data from all 5 cell lines was used as input. Hallmarks and Gene Ontology: Biological Processes gene set collections were used to detect enriched gene sets. Hallmark gene sets enriched in at least 3 out of 5 cell lines (in the same direction, negatively or positively) and with an FDR<0.25 were included, and the top 20 negatively enriched Biological Processes were plotted. Cancer cell line proteomic data was downloaded from the DepMap portal and analyses were performed on the normalised protein quantitation data of 375 cell lines performed as described previously^38^. Proteins which were not detected in at least 50% of the profiled cell lines were removed from the analysis, and cell lines were ordered based on PBRM1 Log2FC. Heatmaps were generated in R using ComplexHeatmap (v2.14.0)^39^ and clustering was performed using default Euclidean distance methods.

### In vivo studies

2 x 10^5^ B16-F10 cells (parental or PBRM1 KOs) were resuspended in 100µL PBS and injected subcutaneously into the right flank of 6-8 week old female C57BL/6J mice (Charles River Laboratories). After 3 days growth, mice were treated with vehicle or 50mg/kg BOS172722 via oral gavage. Mice were treated twice weekly (Monday and Thursday) until the experiment endpoint (Day 42) or were culled once the tumours reached 15mm size in any direction.

### CUT&RUN sequencing

CUT&RUN (Cleavage Under Targets & Release Using Nuclease) was performed according to the CUT&RUN Assay Kit protocol (Cell Signaling Technology) with the following modifications. Cells were detached with Accutase (Sigma). 2×10^5^ cells were collected per experiment and pelleted by centrifugation for 5 minutes at 600 x g. Beads were incubated in the indicated primary antibody (Supplementary Table S2) on a nutator overnight at 4°C. MNase digestion was carried out at 0°C in an ice water bath for 30 minutes. Salt fractionation and DNA purification were carried out according to the CUT&RUN.salt protocol^16^. Briefly, after STOP buffer addition, samples were incubated at 4°C for 1 hour, and the supernatant containing the low salt fraction was collected. Beads were resuspended in low salt buffer (175mM NaCl, 10mM EDTA, 2mM EGTA, 0.1% TritonX-100, 20μg/mL glycogen). High salt buffer (825mM NaCl, 10mM EDTA, 2mM EGTA, 0.1% TritonX-100, 20μg/mL glycogen) was added to beads dropwise with gentle vortexing. Samples were again rocked at 4°C for 1 hour, centrifuged for 5 minutes at 16,000 x g, and the supernatant containing the high salt fraction was collected. Low salt fractions were adjusted to 500mM NaCl. RNAse A (Thermo Fisher Scientific) was added to all fractions and samples were incubated at 37°C for 20 minutes. SDS (sodium dodecyl sulfate) and Proteinase K (Cell Signaling Technology) were added and samples were incubated at 50°C for 1 hour. DNA was extracted using phenol/chloroform and precipitated in 100% ethanol as described before library preparation. Libraries were prepared using the NEBNext Ultra II DNA library prep kit for Illumina (New England Biolabs), profiled using the Agilent TapeStation D1000 high sensitivity ScreenTape on the Agilent 4150 TapeStation System, and sequenced on the Illumina Novaseq 6000 with 150x150 bp reads by Novogene (Novogene Corporation Inc. UK). For PBRM1 knockout cells, replicates 1 and 2 were performed with KO1 and replicate 3 was performed with KO2.

### CUT&RUN analysis

Fastq reads were trimmed using TrimGalore (v0.6.6) (Krueger F, Trimgalore (2023), GitHub repository, https://github.com/FelixKrueger/TrimGalore) using the options --trim-n --paired. The human CHM13-T2Tv1.1 (GenBank GCA_009914755.3) and S. cerevisiae S288C (GenBank GCA_000146045.2) assemblies were combined into one FASTA file, then trimmed reads were mapped using Bowtie2 (v2.4.2)^40^ using the parameters --local --very-sensitive-local --no-unal --no-mixed --no-discordant --dovetail --soft-clipped-unmapped-tlen -- non-deterministic --phred33 -I 50 -X 1500. Reads with more than 3 mismatches were removed with Sambamba (v0.5.0)^41^, and their corresponding mates were removed with Picard tools (v2.23.8) (“Picard Toolkit.” 2019. Broad Institute, GitHub Repository. https://broadinstitute.github.io/picard/; Broad Institute). Sam files were converted to bam with SAMtools (v1.11)^42^. Reads mapped to the CHM13-T2Tv1.1 assembly were extracted using BAMtools (v2.5.1)^43^ split. Duplicate reads were removed using Picard if deemed necessary. Low and high salt BAM files from each sample were merged, sorted, and indexed using SAMtools.

### Uniquely mapped reads analysis

Uniquely mapping reads were extracted with Sambamba^41^, using the parameters: view -h -t 6 -f bam -F "[XS] == null and not unmapped and not duplicate" Bigwig files were made using Deeptools (bamCoverage --extendReads --maxFragmentLength 1500 --binSize 10 – outFileFormat bigwig, v3.1.3)^44^ and normalised by total number of mapped reads, including non-uniquely mapping ones (using --scaleFactor parameter). Peak calling was performed on uniquely mapping read bam files using MACS2 (v2.2.7.1)^45^ callpeak using IgG as control, with options -g 3054832041 -f BAMPE --keep-dup all -q 0.01 --broad --broad-cutoff 0.01.

Bedtools (v 2.29.2)^46^ intersect was used *** . Screenshots of bigwig files and peaks were taken from IGV (v2.11.9)^47^. The ChIPseeker R package (v1.40.0)^48^ was used to analyse the genomic distribution of peaks across features. The intervene package (v0.6.5)^49^ was used to calculate overlapping peak intersections and resulting venn diagrams were re-drawn using the ggVennDiagram R package (v1.5.2)^50^. The EnrichedHeatmap R package (v1.34.0)^51^ was used to generate heatmaps of CUT&RUN and ChIP-seq signal across regions extended +/- 5kb around target peaks, with an averaged signal plot above.

### Analysis of downloaded publicly available ChIP-seq datasets

Histone mark ChIP-seq data for RPE1 cells across the cell cycle, and associated input data was downloaded from GEO accession GSE175752^23^. Fastq files across the cell cycle (G1, ES, LS, G2) in RPE.Ctrl cells were merged for downstream analysis. ChIP-seq data of PBRM1 (SRR12036678, SRR23588908), SMARCA4 (SRR12036684, SRR23588912) and input (SRR12036698) in parental 786-O cells, as well as PBRM1 (SRR12036715, SRR23588903) and input (SRR12036716) from A498, and PBRM1 (SRR12036717-SRR12036722) and input (SRR12036723- SRR12036728) from HK2 were downloaded from the GEO accession GSE152681^20^. Fastq files, where multiple SRA files were available for the same sample, were merged for downstream analysis. ChIP-seq data of SMARCA4 (SRR2133615) and input (SRR2133624) from parental HCT116 cells were downloaded from the GEO accession GSE71510^21^. These ChIP-seq datasets were analysed as above for the CUT&RUN with a few exceptions, as they have single-end reads rather than paired-end reads, the Picard tools step to remove corresponding mates following removing reads with 3 mismatches was no longer required. Bowtie2 parameters were changed to -I 300 -X 700 for analysing GSE152681. Following uniquely mapping read filtering, bigwig files were generated with updated bamCoverage parameters to not include –maxFragmentLength, but to update to --extendReads 250 (for GSE175752), --extendReads 500 (for GSE152681) and --extendReads 180 (for GSE71510).

### Read filtering steps for *k*-mer analysis

The seqkit (v2.5.1)^52^ fx2tab function was used to identify read IDs of CUT&RUN reads that mapped to the spike-in, S. cerevisiae S288C (converted bam file to fastq using bedtools bamtofastq (v 2.29.2)^46^. The BBmap (v38.84) (BBMap – Bushnell B. – sourceforge.net/projects/bbmap/) filterbyname.sh tool was used to filter out and exclude reads mapping to the S. cerevisiae S288C genome from the original fastq files. CUT&RUN reads mapped to the CHM13-T2Tv1.1 assembly, following extraction of S. cerevisiae S288C reads, were used to calculate the total base count for each replicate for normalisation purposes, following deduplication, trimming and interleaving of paired reads (described below). Reads which intersected with the centromere and pericentromere were extracted for full *k*-mer analysis. Briefly, sorted bam files were converted to paired end bed files using bedtools (v 2.29.2)^46^ bamtobed -bedpe function, and then reads that mapped to the centromere and pericentromere were extracted using bedtools intersect function with the CHM13-T2Tv1.1 cenSatAnnotation.bed file. Intersecting read names were printed and used with the BBmap (BBMap – Bushnell B. – sourceforge.net/projects/bbmap/) filterbyname.sh tool to filter to only include centromere and pericentromere mapping reads from the original fastq files. These reads were then used for the full *k*-mer analysis.

### *k*-mer analysis pipeline

The (reference-free) *k*-mer analysis pipeline was adapted from previously described methods^22,1^. First, the clumpify.sh tool from BBmap was used to remove duplicate paired reads that could be PCR duplicates. Then, trimmomatic (v0.39)^53^ was used to trim adapters from paired reads, using the following parameters PE -phred33 ILLUMINACLIP: TruSeq3- PE.fa:2:30:12 SLIDINGWINDOW:10:10 MINLEN:71. The reformat.sh tool from BBmap was used to interleave paired reads for subsequent analysis. Then the kmc package (v3.2.1)^54^ and kmc tools command were used to count and dump 51bp *k*-mers that occur at least twice in each replicate, using the parameters -k51 -sm -ci2 -cs100000000. The R package dplyr (v1.1.0)(Wickham H, François R, Henry L, Müller K, Vaughan D (2023). _dplyr: A Grammar of Data Manipulation_) was used to ‘full_join’ the *k*-mer counts of all replicates and any missing values were replaced with a value of 1 (as missing values could have a count of 0 or 1, so to underestimate enrichment, and to allow fold change calculation). *K*-mer counts were divided by the total base counts for each replicate (from the total genome reads) to calculate the normalised *k*-mer counts. Low and high salt fastq files were analysed separately and their normalised I-mer counts were added together for downstream analysis. *K*-mers were filtered to only include those with a normalized *k*-mer count of >5e-09 for each replicate individually, and then fold changes were calculated, comparing one sample replicate against the averaged IgG normalised *k*-mer count as a control (e.g. SMARCA4 vs average IgG).

Note that IgG replicates were included in the average from each experiment (n=3 for SMARCA4 and CENPB in parental RPE1 cells, n=3 for PBRM1 in parental RPE1 cells, and n=3 for SMARCA4 in PBRM1 knockout cells). Tidyverse (v2.0.0) (doi:10.21105/joss.01686) and tibble (v3.2.0) (Müller K, Wickham H (2022). _tibble: Simple Data Frames_) packages were also used above for data formatting purposes, using r-base v4.2.3.

### Enriched *k*-mer selection and downstream analysis

Enriched *k*-mers were selected if they had a fold change of >2 in the normalised *k*-mer count of at least 2 replicates vs the averaged IgG normalised *k*-mer count. Enriched *k*-mer output files were converted to FASTA format, using sed ’1d’ and awk -F ’[ ]’ ’BEGIN{{OFS="\n"}}{{n=NR; x=">"n; print x, $1}}’. Bowtie2 (v2.4.2)^40^ was used to map these *k*-mers back to the CHM13-T2Tv1.1 genome, using the specific parameters -f -k 5000 -- score-min C,0,0 to allow only exact sequences to be mapped back up to 5000 times.

Bedtools (v 2.29.2)^46^ intersect was again used to extract *k*-mer mapping sites that intersect with the centromere annotation, and awk -F [“_”,”\t”] ‘OFS=”\t” ‘^1^’ was used to split the centromere annotation in the subsequent bed file from e.g. ct_1_1(p_arm) to ct. The GNU datamash package (v1.1.0) (Free Software Foundation, I. (2014). GNU Datamash. Retrieved from https://www.gnu.org/software/datamash/) was then used to create a pivot table summarising how many times each *k*-mer mapped to each part of the centromere annotation (following the split described above), using the following script: sort -k 1 | datamash -s crosstab 1,2 | sed ’s∼N/A∼0∼g’. Bigwig files were generated from enriched k-mer bam files using deepTools bamCoverage tool using the following parameters --smoothLength 1 -- binSize 1 --scaleFactor 1 and screenshots were taken from IGV^47^. Motifs associated with enriched *k*-mers were discovered using the MEME-suite SEA *k*-mer enrichment package^55^ . Venn diagrams were created with the ggVennDiagram R package^50^. Bedtools (v 2.29.2) reldist function^46^ (based on^24^) was used to calculate the relative distance between a query set (*e.g.* H3K9me3 *k*-mer mapping locations) and a reference set (*e.g.* PBRM1-specific *k*- mer mapping locations). This was calculated using the following formula: min(d1,d2)/r – where d1 and d2 are the distances between each location of the query set and the two closest locations of the reference set, and r is the total distance between the two closest locations of the reference set, *i.e.* (d1+d2). The cumulative fractions of query *k*-mer mapping locations at each relative distance up to a maximum possible of 0.5 were calculated and plotted.

### RNA-sequencing

For RNA-sequencing, pellets were harvested by scraping cells in ice-cold PBS following by centrifugation at 7,500 x g for 10 minutes. Three independent biological replicates were used for parental cells (n=3), and two independent biological replicates of two individual PBRM1 knockout clones were used for PBRM1 knockouts, which were combined for n=4 in total.

Pellets were resuspended in 500µL TRIzol reagent and total RNA was extracted using a Direct-zol RNA miniprep kit (Zymo Research) according to the manufacturer’s protocol. RNA concentration and quality was confirmed with the Agilent High Sensitivity RNA ScreenTape, using the Agilent Tapestation as above. Library preparation and sequencing was performed by Novogene Corporation Ltd. Novogene NGS Stranded RNA Library Prep Set was used to generate 250-300bp insert strand specific libraries, and ribosomal RNA was removed using TruSeq Stranded Total RNA Library Prep. 50 million 150bp paired-end reads were sequenced on an Illumina NovaSeq 6000. Fastq reads were checked using FastQC (v0.11.9) (Andrews, S. (2010). FastQC: A Quality Control Tool for High Throughput Sequence Data [Online]. http://www.bioinformatics.babraham.ac.uk/projects/fastqc/) and trimmed using TrimGalore (v0.6.6) (Krueger F, Trimgalore (2023), GitHub repository, https://github.com/FelixKrueger/TrimGalore). Residual ribosomal RNA reads were removed using Ribodetector with -e norRNA setting (v0.2.7)^56^ and strandedness was detected using RSeQC (v4.0.0)^57^. Reads were aligned to the T2T-CHM13v2 genome using STAR alignment software (v2.7.6a)^58^ and reads mapping to genes were quantified using HTSeqCount (v0.12.4)^59^. Differential analysis of gene expression was calculated in R using DESeq2 (v1.38.3)^60^.

## Software & statistical analyses

The number of biological repeats for each experiment and statistical analyses used are indicated in the Fig. legend for each experiment. Graphs were generated and statistical analyses performed using GraphPad Prism (v9.5.1). Outliers were removed using Grubbs’ test using a significance level of 0.05. Microscopy images were analysed using ImageJ (v1.5.3 or 1.5.4) or CellProfiler (v4.0.7) software, and visualized using GraphPad Prism or ggplot2 (v3.4.4)(Valero-Mora (2010). ggplot2: Elegant Graphics for Data Analysis. https://doi.org/10.18637/jss.v035.b01). For statistical analyses of microscopy data, the median of each biological repeat was used for the appropriate statistical test, described in the corresponding Fig. legend). Omics data were visualised using the indicated packages in R (v4.2.1, 4.2.2, or 4.4.0 except where noted) and R Studio (v2021.09.0, 2023.03.1, 2024.04.2 except where noted). Fig.s were generated using Adobe Illustrator (v27.5). Some graphics were generated using BioRender (biorender.com).

## Acknowledgements

We thank the Institute of Cancer Research Flow Cytometry, Light Microscopy and the Biological Services Unit for support, and we thank the ICR Scientific Computing Team for HPC services. We thank all members of the Downs lab, Jon Pines, and Bill Earnshaw for helpful discussions, Swen Hoelder and Florence Raynaud for reagents and advice, and former lab members Peter Brownlee and Cornelia Meisenberg for early observations. This work was supported by Cancer Research UK (C7905/A25715, C309/A25144, and DRCRPGTD- Nov21100001) and the Medical Research Council (MR/W001276/1).

## Author contributions

KAL, AH, and JAD conceived the project. KAL, AH, LW, TIR, HF, SF, KB, and FS designed and performed experiments. AH, LW, and TIR performed bioinformatics analyses. FTZ, AAM, JCS and JAD obtained funding and provided input and supervision. KAL, AH, and JAD prepared the original draft, and all authors contributed to review and editing.

## Data availability

The datasets generated in this study are available in the GEO repository (accession numbers GSE235342 and GSE235294) or ProteomeXchange Consortium via the PRIDE repository (PXD043209). All cell lines used in this study have been made available at CancerTools.org. All data generated in this study are provided in the Supplementary Information figures and tables.

---

**Figure S1.**
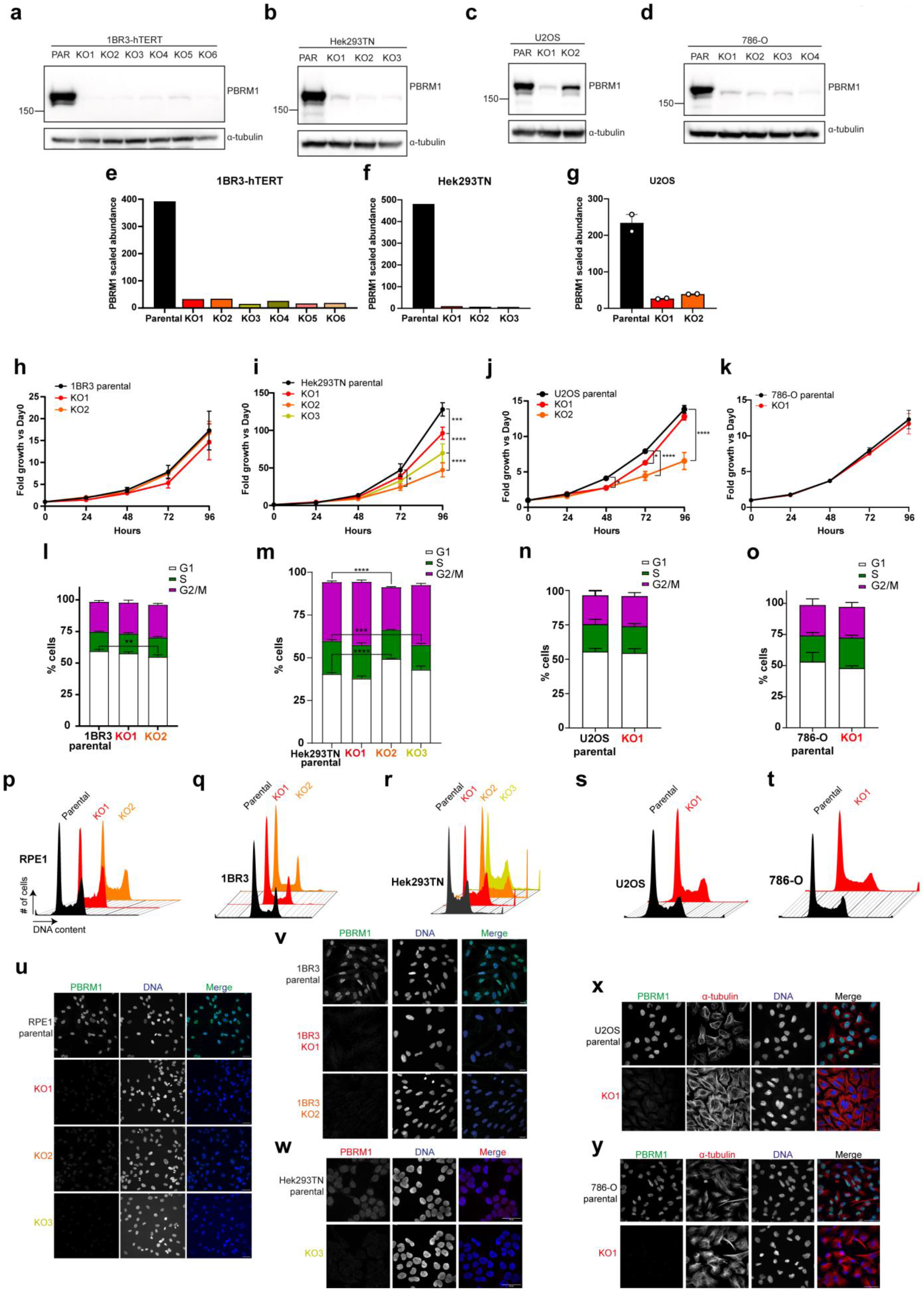
Characterisation of PBRM1 KO cell lines. **(a)-(d)** Western blotting for PBRM1 in parental and PBRM1 knockouts of the indicated cell lines – (a) 1BR3-hTERT, (b) Hek293TN, (c) U2OS, and (d) 786-O. Note that one clone in U2OS (“HetKO2”) retains low levels of PBRM1 due to a heterozygous truncating mutation. α-tubulin was used as loading control. **(e)-(g)** Scaled abundances of PBRM1 in proteomic analyses of whole cell protein extracts of parental and PBRM1 knockouts in (e) 1BR3-hTERT, (f) Hek293TN, and (g) U2OS. Points in (g) correspond to independent biological replicates (*n*=2). PBRM1 was not detected in proteomic analyses of 786-O cells. **(h)-(k)** Proliferation rates of PBRM1 knockouts in (h) 1BR3-hTERT, (i) Hek293TN, (j) U2OS, and (k) 786-O, compared to parental cells. Cells were counted every 24 hours and fold change in cell number was calculated compared to the number of cells seeded. *n*=2- 4, mean±SEM, and data were analysed by 2way ANOVA with Dunnett’s test, \*\**p*<0.01, \*\*\**p*<0.001, \*\*\*\**p*<0.0001. **(l)-(o)** Cell cycle distribution of parental and PBRM1 knockout cells in (l) 1BR3-hTERT, (m) Hek293TN, (n) U2OS, and (o) 786-O, measured using flow cytometry. *n*=3-4, mean±SEM and data were analysed by 2way ANOVA with Dunnett’s test, \**p*<0.05, \**p*<0.001, \*\*\*\**p*<0.0001. **(p)-(t)** Representative histograms showing cell cycle profiles of parental and PBRM1 knockout (p) hTERT-RPE1, (q) 1BR3-hTERT, (r) Hek293TN, (s) U2OS, and (t) 786-O cells, corresponding to data shown in Figure 1(e) and S1(l)-(o). **(u)-(y)** Representative immunofluorescence images showing PBRM1 expression in parental and PBRM1 knockout cells in (u) hTERT-RPE1, (v) 1BR3-hTERT, (w) Hek293TN, (x) U2OS, and (y) 786-O, as well as nuclear (DNA, blue) staining. (x) & (y) also show cytoskeleton (α-tubulin, red) staining. Scale bars correspond to 20µm in (u), (v), & (x), and correspond to 50 µM in (w) and (y).

**Figure S2.**
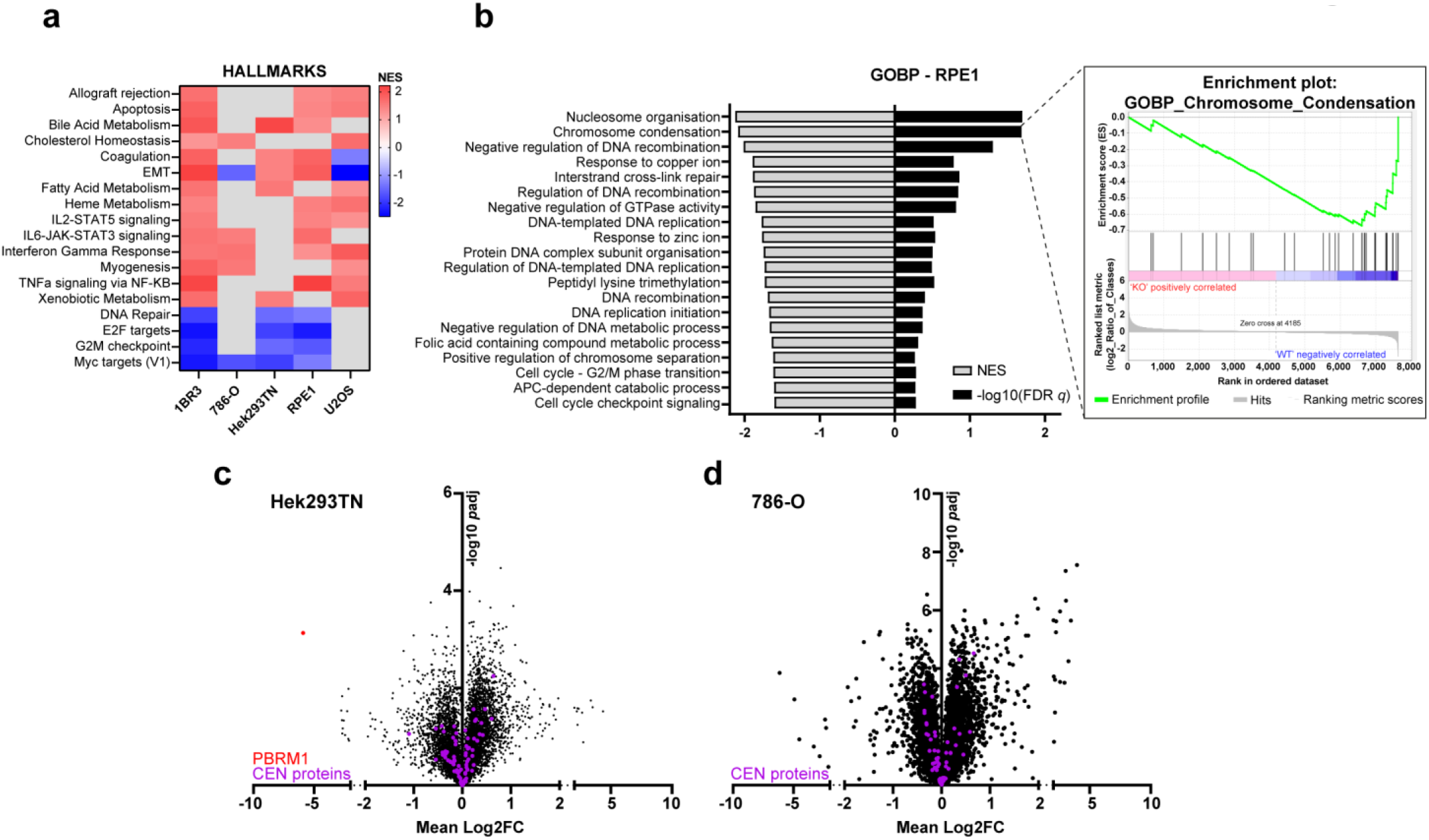
Dysregulation of a number of pathways is conserved across PBRM1 KO cell lines. **(a)** Heatmap showing the normalised enrichment score (NES) of Hallmark gene sets associated with PBRM1 loss, determined using GSEA analysis. Red corresponds to positive enrichment in PBRM1 knockouts versus the corresponding parental cells, while blue corresponds to negatively enriched gene sets. Grey cells indicate gene sets which did not reach a significant enrichment level of FDR <0.25. Only Hallmark gene sets significantly enriched in 3 or more cell lines were included on heatmap. **(b)** GSEA gene ontology analysis of biological processes (GOBP) negatively enriched in RPE1 PBRM1 knockouts compared to parental cells. The 20 processes with the most significant FDR *q* values were included. NES (grey bars) of these enriched biological processes, as well as the -log10 FDR *q*-value (black bars) is shown. A representative enrichment plot of one of these biological processes, ‘Chromosome Condensation’, is shown on the right. **(c)-(d)** Protein abundances in PBRM1 knockouts compared to parental cells in (c) Hek293TN and (d) 786-O cell lines, detected using LC-MS of whole cell protein extracts. The mean Log2FC of protein abundance in PBRM1 knockouts versus parental cells is plotted against the -log10p. PBRM1 is highlighted in red, while peri/centromeric proteins are highlighted in purple.

**Figure S3.**
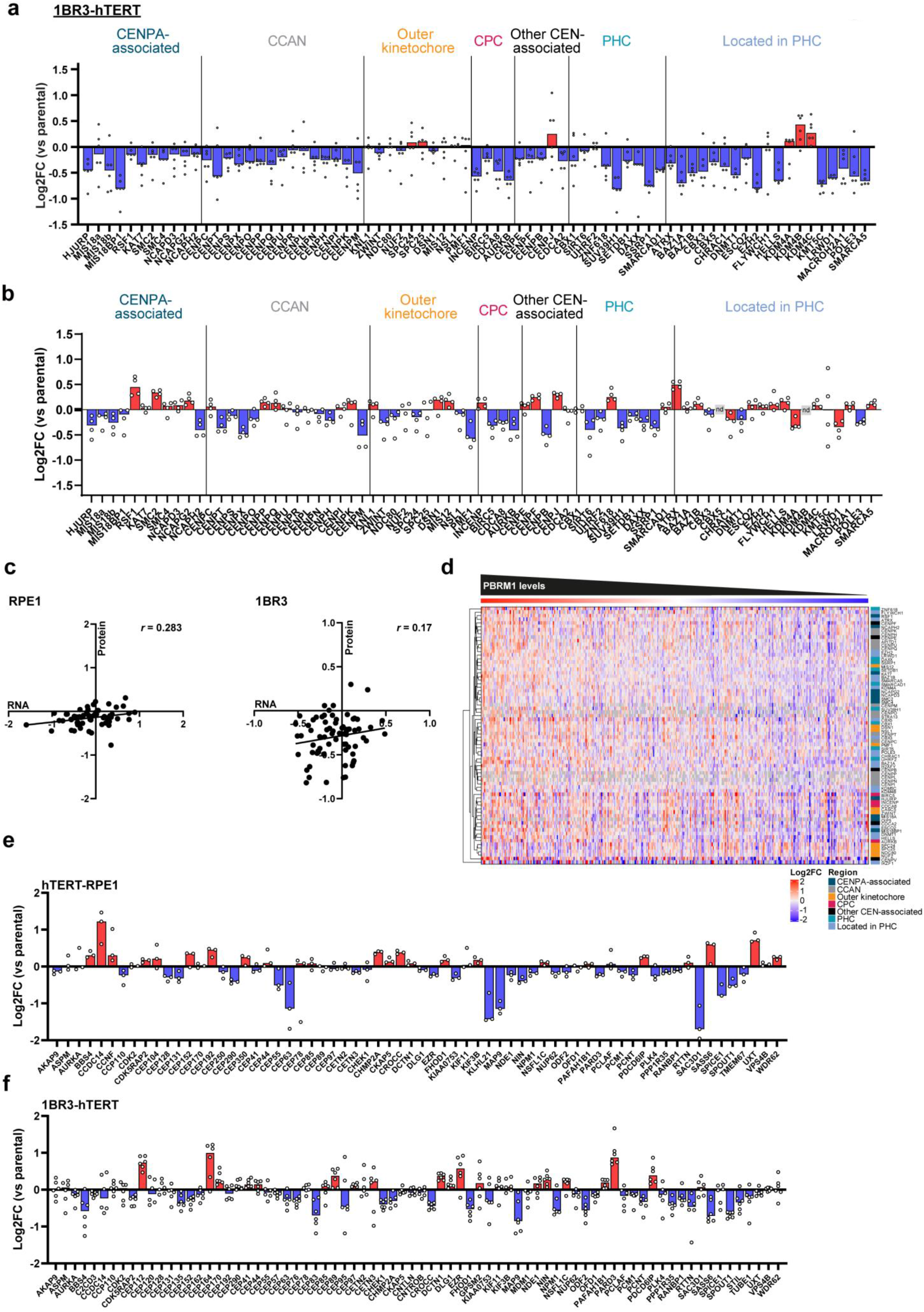
PBRM1 KO cells show a loss of centromere proteins. **(a)** Log2FC of annotated centromere- and pericentromere-associated proteins in 1BR3 PBRM1 knockouts compared to parental cells. Points correspond to independent knockout clones. **(b)** Transcript levels of annotated centromere- & pericentromere-associated genes corresponding to the proteins in (a) were detected using RNA-seq. Median Log2FC of annotated genes transcribing centromere- & pericentromere-associated proteins in RPE1 PBRM1 knockouts was plotted compared to parental cells. Points correspond to two individual knockout clones (KO1 and KO2) and two independent biological replicates. **(c)** Correlation plot showing Pearson correlation between protein and RNA levels of centromere & pericentromere protein/gene Log2FCs, in RPE1 (left) and 1BR3 (right) PBRM1 KOs compared to parental cells. Line indicates simple linear regression and Pearson correlation coefficient (*r*) is displayed. **(d)** Log2 fold changes of the indicated peri/centromere proteins in cancer cell lines from the CCLE database. Heatmap is ordered from highest to lowest PBRM1 Log2FC, left to right **(e)-(f)** Log2FC of annotated centrosome component and associated proteins in PBRM1 knockouts in (e) RPE1 and (f) 1BR3 cells compared to the parental of each cell line. Individual points represent independent PBRM1 knockout clones.

**Figure S4.**
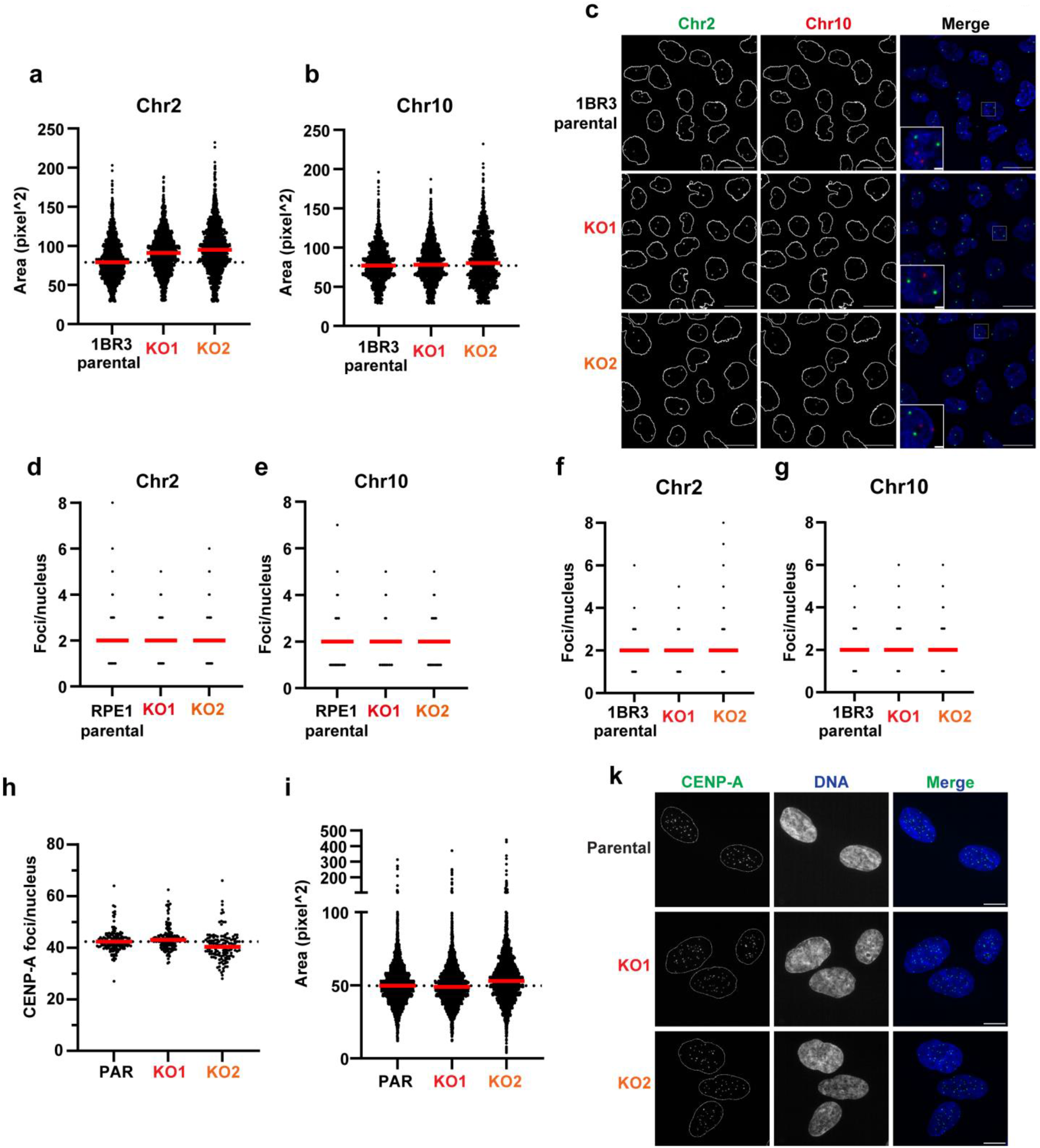
(a)-(b) Quantification of the area of individual foci in 1BR3 parental or PBRM1 knockout cells stained for α- satellite centromeric regions in (a) chromosome 2 and (b) chromosome 10, using FISH probes against α-satellite sequences in the corresponding chromosomes. *n*=3, line at median, and at least 1200 cells were analysed for each cell line. **(c)** Representative images of α-satellite FISH of chromosomes 2 and 10 in 1BR3 parental and PBRM1 knockouts. Scale bars corresponds to 20µm; in zoomed inset images, scale bars correspond to 5µm. **(d)-(e)** Quantification of the number of foci in RPE1 parental or PBRM1 knockout cells stained for α-satellite centromeric regions in (d) chromosome 2 and (e) chromosome 10, using FISH probes against α-satellite sequences in the corresponding chromosomes. *n*=3, line at median. **(f)-(g)** Quantification of the number of foci in 1BR3 parental or PBRM1 knockout cells stained for α-satellite centromeric regions in (f) chromosome 2 and (g) chromosome 10, using FISH probes against α-satellite sequences in the corresponding chromosomes. *n*=3, line at median. **(h)** Quantification of the number of CENP-A foci per nucleus in RPE1 parental and PBRM1 KO cells. *n*=3, line at median, and data were analysed by 1way ANOVA with Dunnett’s test. At least 300 cells were analysed for each condition. **(i)** Quantification of the area of CENP-A foci in RPE1 parental and PBRM1 KO cells. *n*=3, line at median, and data were analysed by 1way ANOVA with Dunnett’s test. At least 300 cells were analysed for each condition. **(j)** Representative images showing CENP-A intensity (green) in RPE1 parental and PBRM1 knockout cells. Scale bars correspond to 20µm.

**Figure S5.**
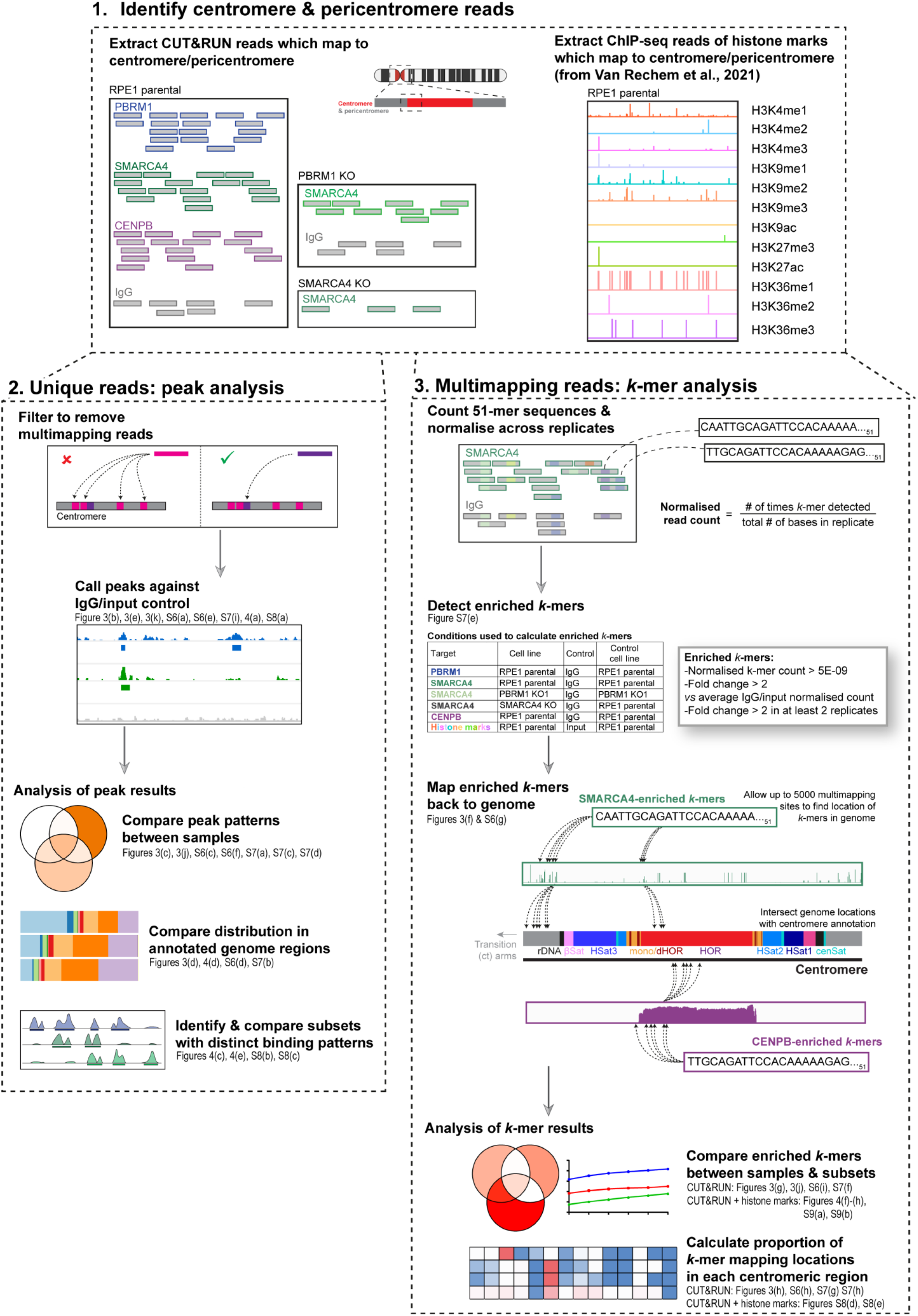
Detailed workflow for the mapping strategy of CUT&RUN and ChIP-seq reads to centromeric and pericentromeric sequences. Workflow for mapping SWI/SNF subunit and CENP-B CUT&RUN and publicly available ChIP-seq reads to centromeric and pericentromeric sequences (Box 1), either by calling peaks only on uniquely mapping reads (Box 2), or by taking all reads that mapped to the centromere and pericentromere, including multimapping reads at repetitive sequences, using *k*-mer enrichment analysis (Box 3). Detailed steps are described in Methods. Briefly, enriched peaks of uniquely mapping reads were compared between conditions and distribution of peaks among features was determined. For enriched *k*-mer analysis, sequencing reads which mapped to the centromeric and pericentromeric regions were extracted and 51-mer sequences in these reads were counted, normalised and compared across samples. Enriched *k*-mers were compared between conditions and were also mapped back to the genome and intersected with centromere annotation to determine where enriched k-mers are located. Data corresponding to these analyses are presented in Figures 3 and 4, and S6-S9, as indicated beside each stage of the analysis workflow.

**Figure S6.**
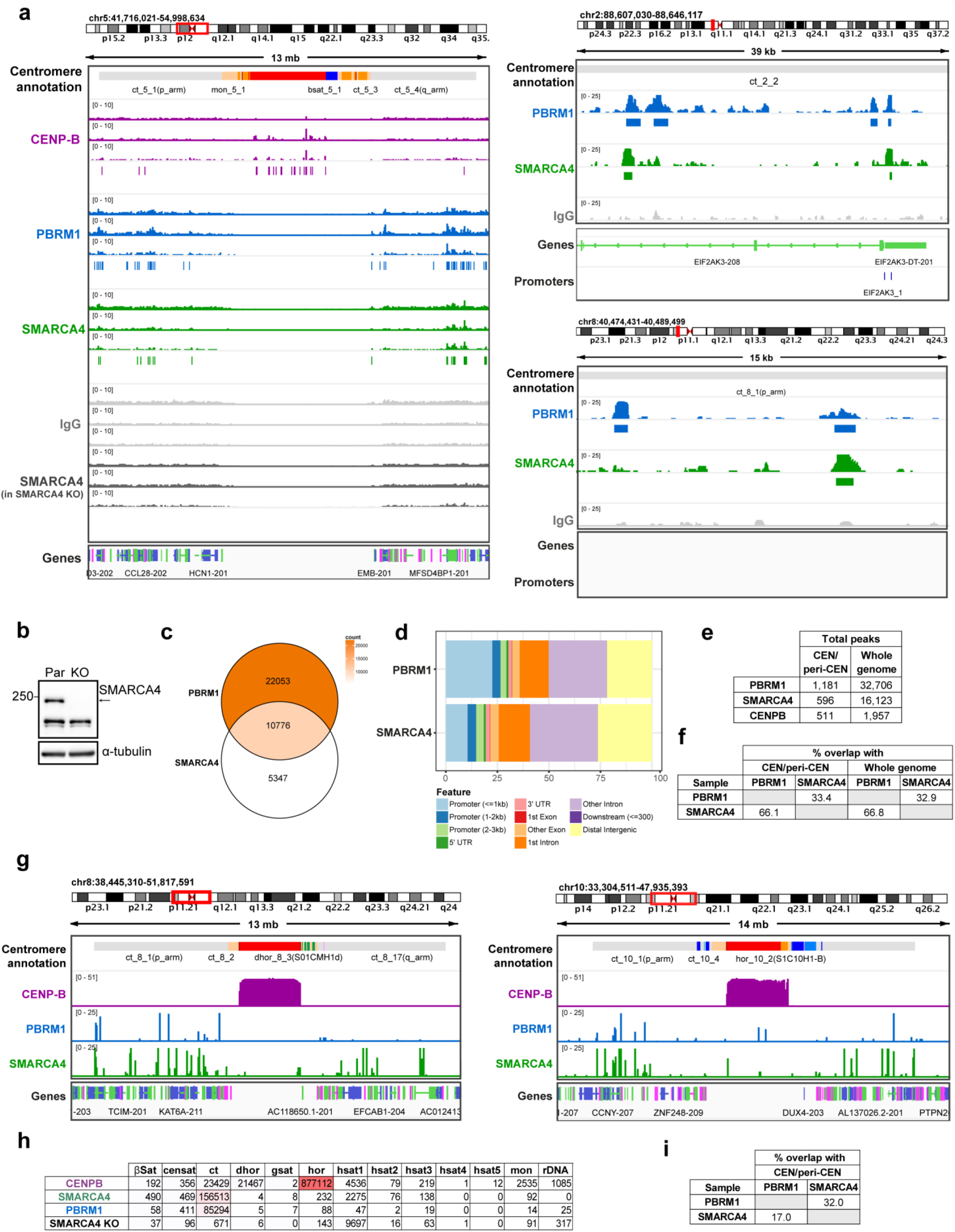
SWI/SNF binding in centromeric and pericentromeric regions. **(a)** Additional examples of representative genome tracks displaying coverage of reads from CENP-B (purple), PBRM1 (blue), SMARCA4 (green) and IgG control (grey) CUT&RUN sequencing in RPE1 parental cells. The full centromere is shown (left), and two examples of a zoomed in view of the centromere transition (ct) arms (right), excluding the CENP-B track. **(b)** Western blotting for SMARCA4 expression in whole cell lysates from RPE1 parental and SMARCA4 knockout cells used in CUT&RUN experiments. α-tubulin is used as loading control. **(c)** Venn diagram indicating the overlap of significantly enriched peaks in the whole genome, in SMARCA4 and PBRM1 in RPE1 parental cells versus their IgG control. Venn diagram shows enriched peaks found in at least two out of the three independent biological replicates, versus their IgG controls. The colour corresponds to the total number of enriched peaks in each region of the Venn diagram (count). **(d)** Stacked colour bar representing the genomic distribution of enriched PBRM1 and SMARCA4 peaks, categorised by feature across the whole genome. **(e)** Table showing the total number of enriched peaks in PBRM1, SMARCA4, and CENP-B CUT&RUN sequencing in RPE1 parental cells, both in the centromere and pericentromere (‘CEN/peri-CEN’) and in the whole genome. **(f)** Table with percentage overlaps of peaks from the Venn diagrams in Figure 3(c) and S6(c), showing shared peaks in the centromere/pericentromere and whole genome. **(g)** Additional representative genome tracks displaying mapping locations of enriched *k*-mers across the centromere and pericentromere from analysis of CENP-B (purple), PBRM1 (blue) and SMARCA4 (green) CUT&RUN sequencing in RPE1 parental cells). Genomic location is indicated at the top. Centromeric region is indicated above the tracks and transcript annotation is shown below the tracks. **(h)** Numbers of enriched *k*-mers which could map to the indicated region of the centromere and pericentromere in each dataset. **(i)** Table with percentage overlaps of *k*-mers from the Venn diagram in Figure 3(g).

**Figure S7.**
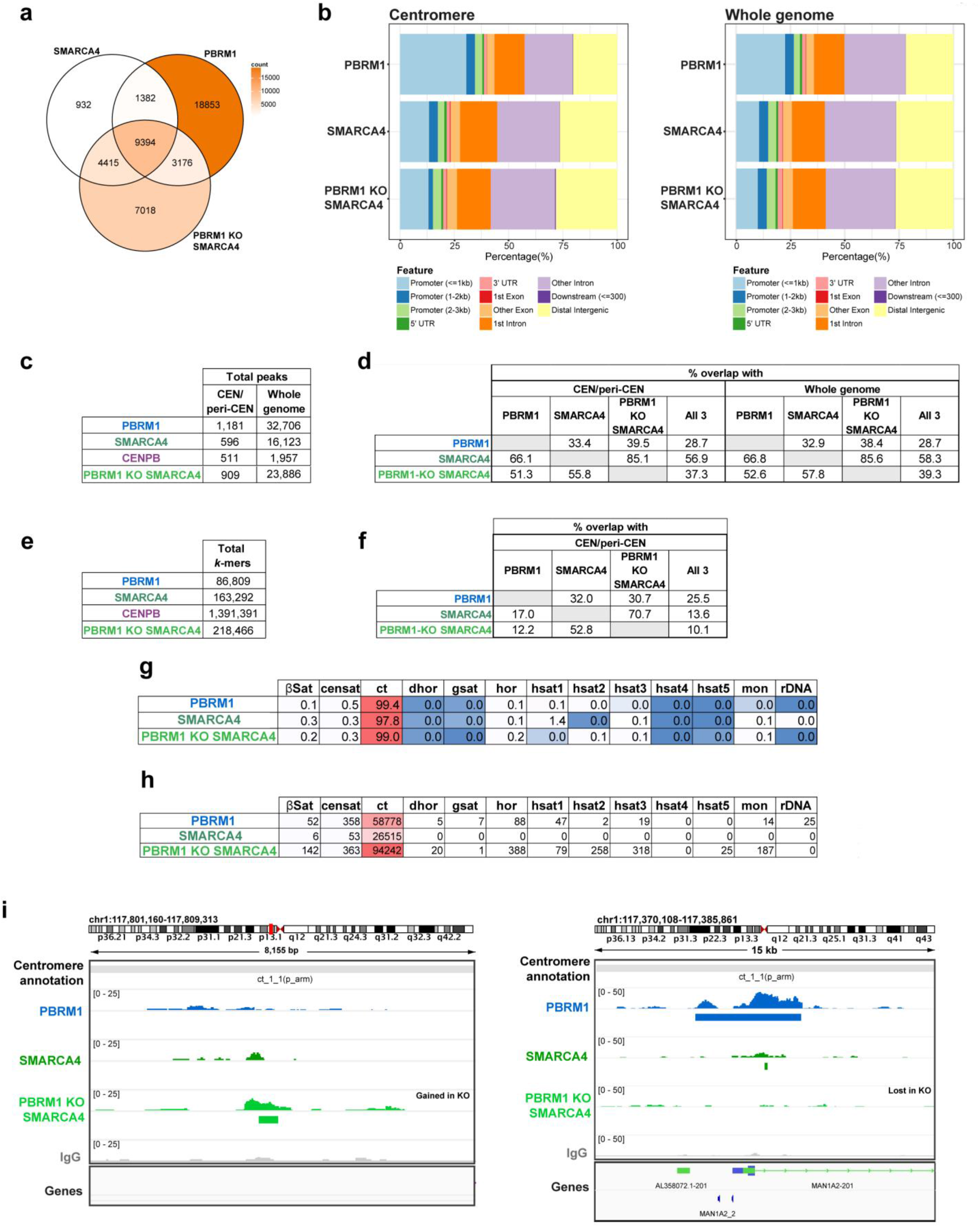
SWI/SNF binding in PBRM1 KO cells is altered. **(a)** Venn diagram indicating the overlap of significantly enriched peaks across the whole genome, in SMARCA4 and PBRM1 in RPE1 parental cells (as per Figure S6(c)), and SMARCA4 in PBRM1 KO cells. Venn diagrams shows enriched peaks found in at least two out of the three independent biological replicates, versus their IgG controls. The colour corresponds to the total number of enriched peaks in each region of the Venn diagram (count). **(b)** Stacked colour bars representing the genomic distribution of enriched PBRM1 and SMARCA4 peaks in RPE1 parental cells (as per Figure 3(d) and Figure S6(d)) and SMARCA4 peaks in PBRM1 knockout cells, categorised by feature, in the centromere and pericentromere (left) and across the whole genome (right). **(c)** Table showing the total number of enriched PBRM1, SMARCA4, and CENP-B peaks in RPE1 parental cells (as per Figure S6(e)), plus SMARCA4 peaks in PBRM1 KO cells, both in the centromere and pericentromere, and in the whole genome. **(d)** Table showing the % overlap of peaks called in PBRM1 and SMARCA4 CUT&RUN sequencing in RPE1 parental cells (as per Figure S6(f)) as well as with SMARCA4 peaks enriched in PBRM1 KO cells, corresponding to Venn diagrams in Figures 3(j) and S7(a). **(e)** Table showing the total number of enriched PBRM1, SMARCA4, and CENP-B *k*-mers in RPE1 parental cells plus SMARCA4 in PBRM1 KO cells, in the centromere and pericentromere. **(f)** Table showing the % overlap of enriched *k*-mers in PBRM1 and SMARCA4 CUT&RUN sequencing in RPE1 parental cells (as per Figure S6(i)) as well as with enriched *k*-mers in SMARCA4 in PBRM1 KO cells, corresponding to Venn diagram in Figures 3(j). **(g)-(h)** Tables displaying (g) percentages and (h) numbers of enriched *k*-mers which could map to the indicated region of the centromere and pericentromere in each condition – PBRM1 and SMARCA4 in RPE1 parental cells (as per Figures 3(h) and S6(h)), as well as SMARCA4 in RPE1 PBRM1 knockout cells. **(i)** Additional representative genome tracks displaying coverage of reads from PBRM1 (blue), SMARCA4 (green) and IgG control (grey) CUT&RUN sequencing in RPE1 parental cells and SMARCA4 in PBRM1 knockout cells (light green), showing an example of peaks gained (left) or lost (right) in PBRM1 knockout cells. One representative independent biological replicate is shown, with boxes underneath representing peaks that were called as significantly enriched (*q*<0.01) in at least two out of the three replicates versus their IgG control. Genomic location is indicated at the top. Centromeric region is indicated above the tracks and transcript annotation is shown below the tracks.

**Figure S8.**
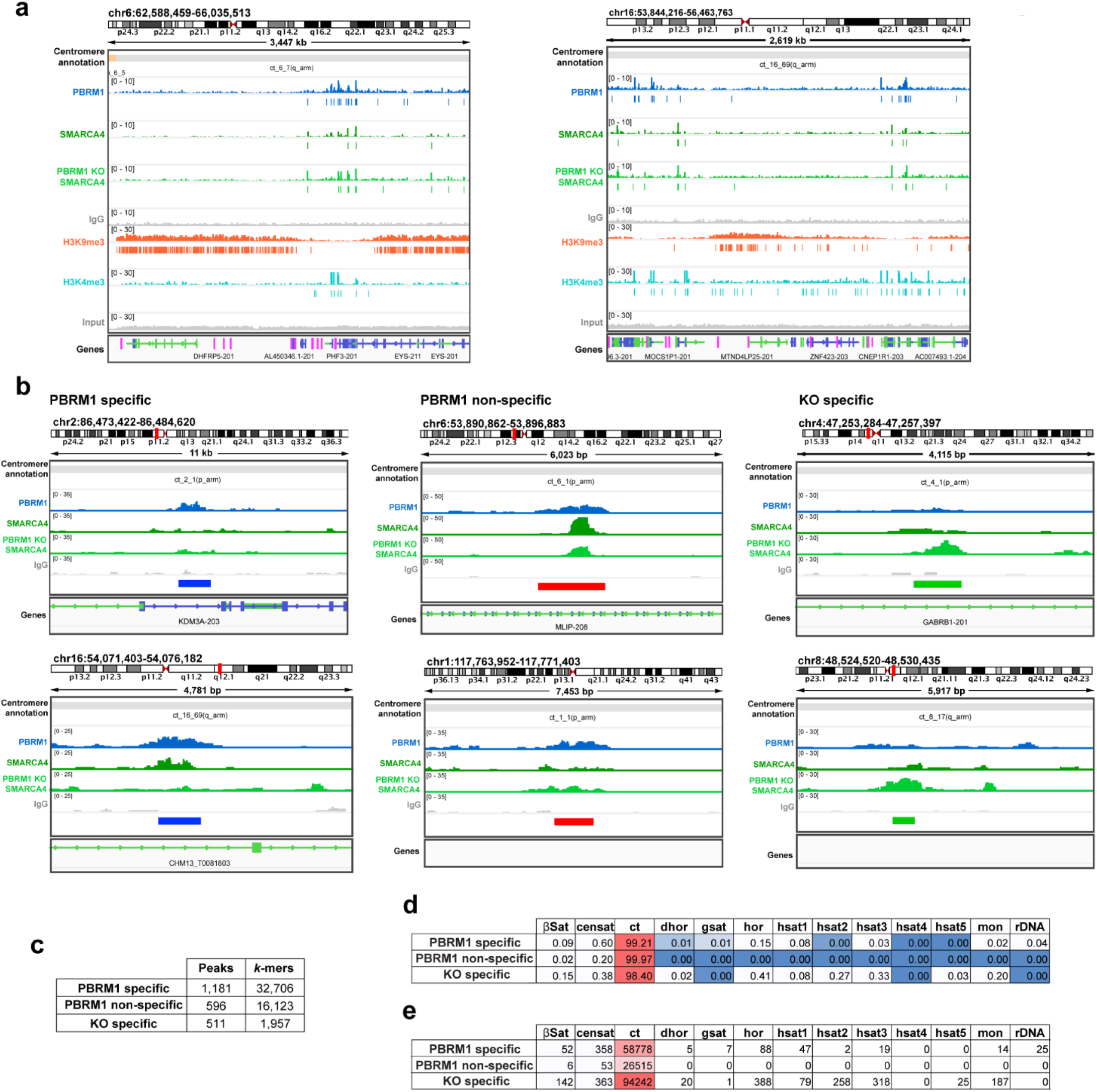
SWI/SNF binding pattern in the centromere and pericentromere changes in the absence of PBRM1 at sites of enriched histone marks in parental RPE1 cells. **(a)** Additional representative genome tracks displaying coverage of reads from PBRM1 (blue), SMARCA4 (green) and IgG control (grey) CUT&RUN sequencing in RPE1 parental cells, SMARCA4 in PBRM1 knockout cells (light green), and H3K9me3 (orange) and H3K4me3 (teal) ChIP-seq in RPE1 parental cells, showing the boundary of heterochromatin in the pericentromere. One representative independent biological replicate is shown, with boxes underneath representing peaks that were called as significantly enriched (*q*<0.01) in at least two replicates versus their IgG or input control. **(b)** Representative genome tracks displaying coverage of reads from PBRM1 (blue), SMARCA4 (green) and IgG control (grey) CUT&RUN sequencing in RPE1 parental cells and SMARCA4 in PBRM1 knockout cells (light green) that highlight the three subsets of peaks (identified as shown in Figure 4(b)), represented as boxes below genome tracks: PBRM1 specific (blue, left), PBRM1 non-specific (red, middle) and KO specific (green, right). For genome browser tracks (a, b), genomic location is indicated at the top, centromeric annotation is shown above the tracks and transcript annotation is shown below. **(c)** Table showing the total number of enriched peaks and *k*-mers from the three subsets of peaks/*k*-mers (PBRM1 specific, PBRM1 non-specific, and KO specific), in the centromere and pericentromere. Subsets of *k*-mers were identified in the same way as peaks (shown in Figure 4(b)). **(d)-(e)** Tables displaying (d) percentages and (e) numbers of enriched *k*-mers which could map to the indicated region of the centromere and pericentromere in each subset of *k*- mers; PBRM1 specific, PBRM1 non-specific and KO-specific.

**Figure S9.**
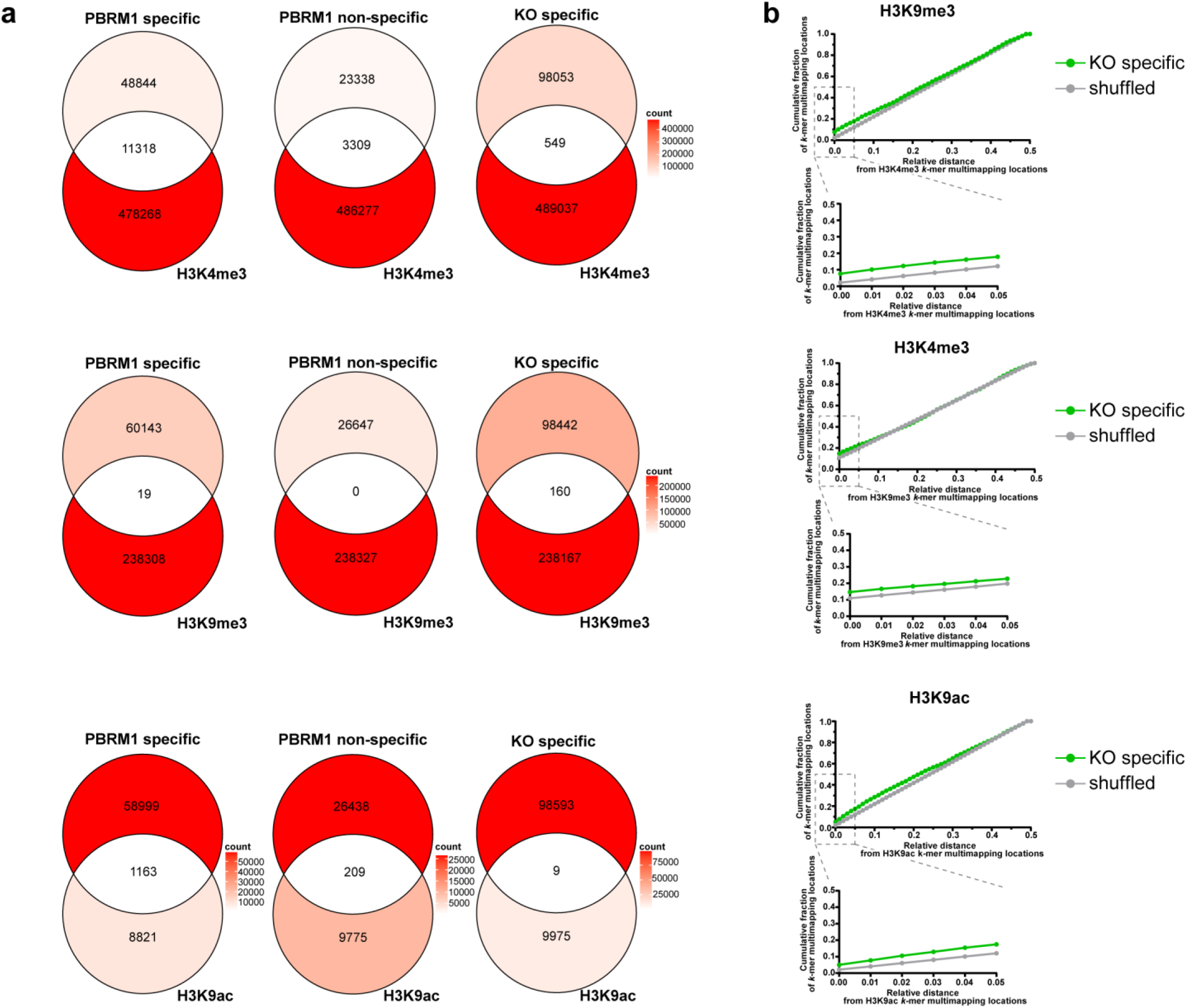
PBAF association with H3K9 modifications is altered in PBRM1 knockouts. **(a)** Venn diagrams indicating the overlap of enriched *k*-mer subsets (PBRM1 specific, left; PBRM1 non-specific, middle; KO-specific, right) in the centromere and pericentromere, with enriched H3K4me3 (top), H3K9me3 (middle) and H3K9ac *k*-mers (bottom). The colour corresponds to the total number of enriched *k*-mers in each region of the Venn diagram (count). **(b)** Plots displaying the cumulative fraction of reference *k*-mer multimapping locations (KO specific (green) and as a control (grey), shuffled locations of KO specific *k*-mer multimapping locations within the centromere and pericentromere at a relative distance from query histone mark *k*-mer multimapping locations (H3K4me3, top; H3K9me3, middle; H3K9ac, bottom). Relative distances are calculated as the shortest distance between each *k*-mer multimapping location in the query set and the two closest *k*-mer multimapping locations in the reference set, divided by the total distance between the two closest *k*-mer multimapping locations, so the maximum relative distance is 0.5. A zoomed in view of the smallest relative distances (0 to 0.05) is shown below each plot.

**Figure S10.**
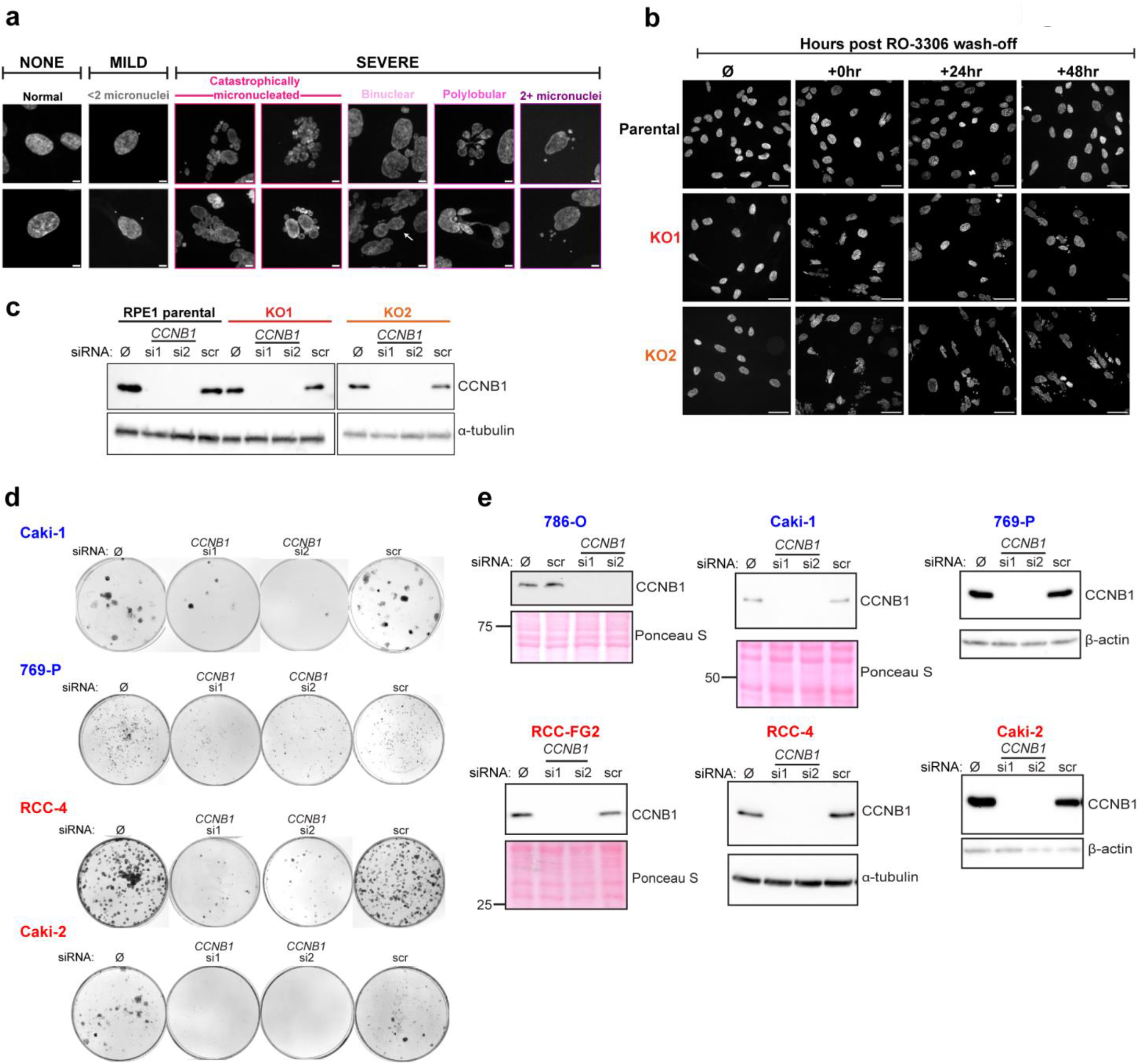
PBRM1-deficient cells are sensitive to mitotic perturbation. **(a)** Representative images of nuclear defects and their associated morphological phenotypes, as quantified in Figure 5(c)-(f). Scale bar represents 5µm. **(B)** Representative images from full timecourse in Figure 5(f), where nuclear defects in RPE1 parental and PBRM1 knockout cells were quantified after treatment with 3µM RO-3306 for 24 hours (or treatment with DMSO alone (Ø)), and allowed to recover for 0hr, 24hr, or 48hr. Scale bar corresponds to 40µm. **(c)** Representative western blot for CCNB1 following CCNB1 RNAi in RPE1 parental and PBRM1 knockout cells, corresponding to clonogenic survival assays in Figure 5(g). **(d)** Representative images of colony formation following CCNB1 RNAi in a panel of PBRM1- proficient (blue) and -deficient (red) RCC cell lines, corresponding to data in Figure 5(i). Representative images from 786-O and RCC-FG2 cell lines are shown in Figure 5(k). **(e)** Representative western blots to detect CCNB1 protein expression after CCNB1 RNAi in a panel of PBRM1-proficient (blue) and -deficient (red) RCC cell lines, corresponding to data in Figure 5(i).

**Figure S11.**
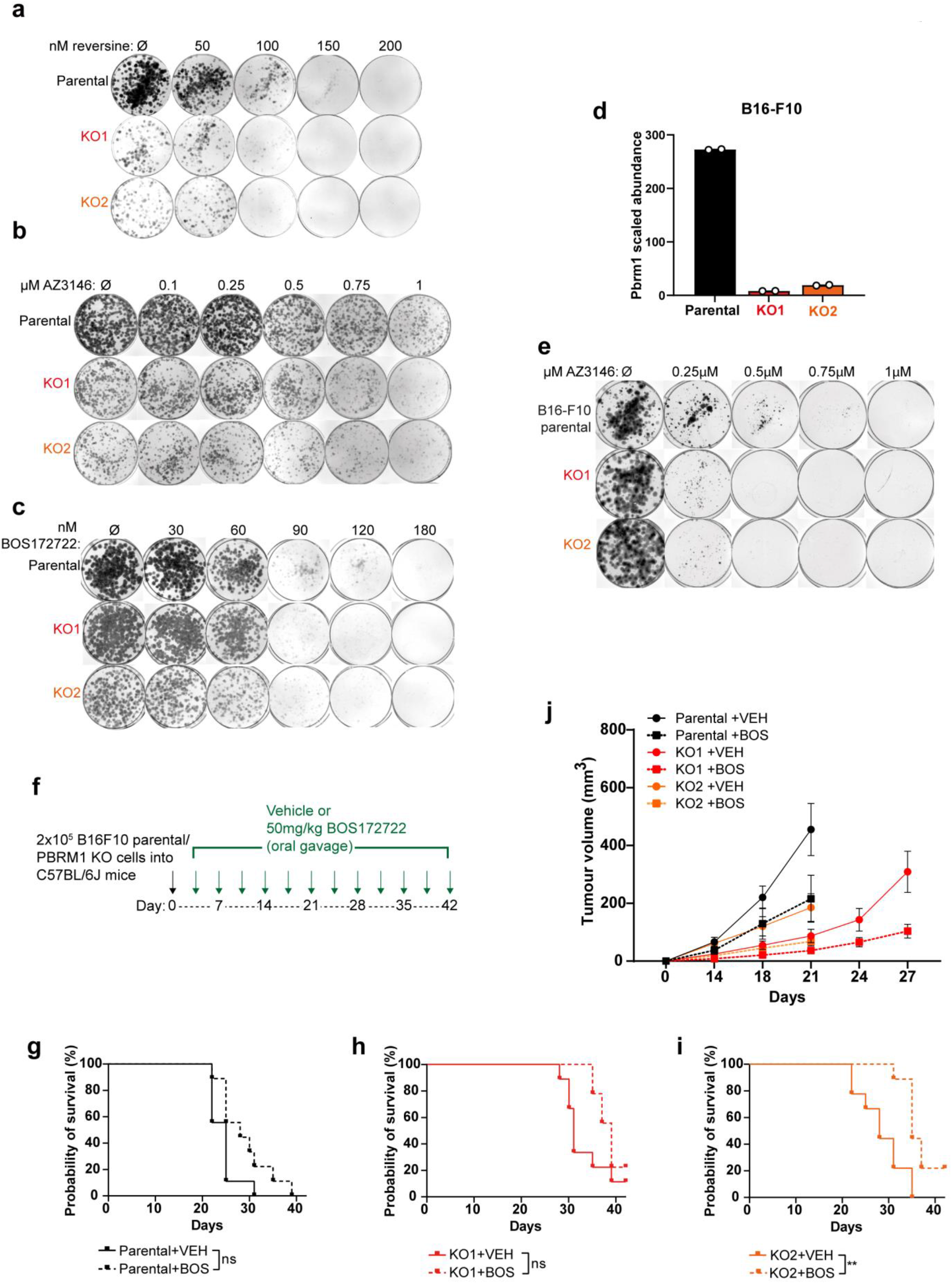
PBRM1 knockouts are sensitive to Mps1 inhibition *in vitro* and *in vivo*. **(a)-(c)** Representative images of colony formation from clonogenic survival assays in Figure 6(a)-(c), comparing survival of RPE1 parental and PBRM1 knockout cells to doses increasing doses of Mps1 inhibitors, compared to colony formation in DMSO-only treated cells (Ø). **(d)** Scaled abundances of Pbrm1 following LC-MS of whole cell protein extracts of B16-F10 parental and PBRM1 knockout cells. Points correspond to independent biological replicates (*n*=2). **(e)** Representative images of colony formation from clonogenic survival assays in Figure 6(f), comparing survival of B16-F10 parental and PBRM1 knockout cells treated with increasing doses of the Mps1 inhibitor AZ3146, compared to colony formation in DMSO-only treated cells (Ø). **(f)** Treatment outline for in vivo studies. C57BL/6J mice were injected with B16-F10 parental or PBRM1 knockout cells (black arrow). After 3 days tumour growth and twice weekly after this, mice were treated with either vehicle or 50mg/kg BOS172722 and their tumour volume was measured (green arrows). **(g)-(i)** Individual Kaplan-Meier survival curves showing survival of mice with tumours derived from B16-F10 (g) parental or (h)-(i) PBRM1 knockouts, treated with vehicle (VEH) or BOS172722 (BOS). *n*=9 mice per condition, and survival was compared using the logrank (Mantel-Cox) test, \*\**p*<0.01. **(j)** Graph showing tumour volumes over indicated times of mice injected with B16-F10 parental or PBRM1 KO cells, treated with either vehicle (VEH) or BOS172722 (BOS). *n*=9 per condition, and graph shows data up until latest timepoint where all mice were still surviving. Tumour growth corresponds to area under the curve (A.U.C.) data from Figures 6(j)-(l). Line indicates mean tumour volume ± SEM.

**Figure S12.**
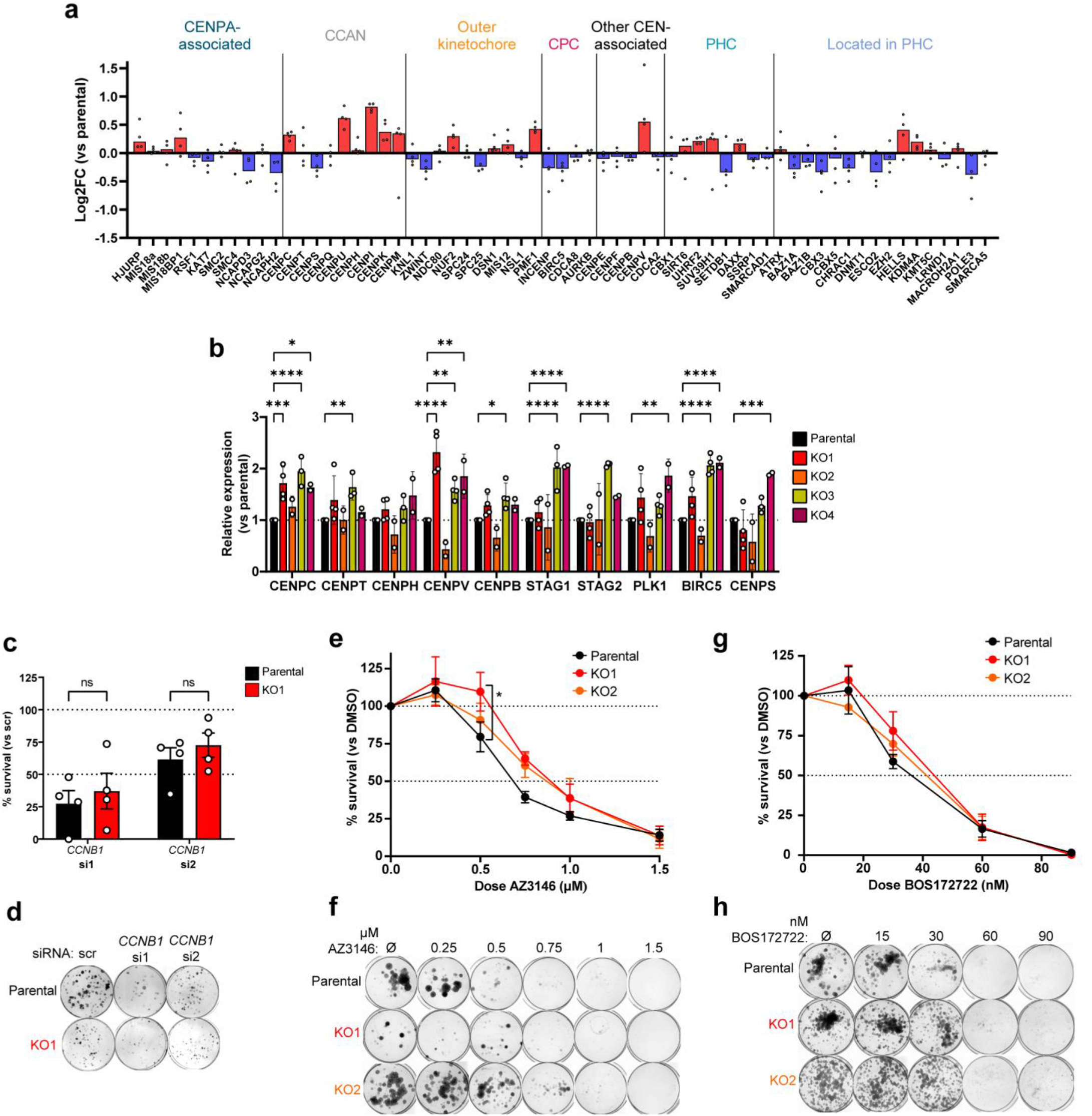
PBRM1 KOs in 786-O do not show exhibit centromere phenotypes. **(a)** Median Log2FC of annotated centromere and pericentromere proteins in RPE1 PBRM1 knockouts compared to parental cells. Points correspond to individual knockout clones from one of two independent biological replicates. **(b)** Expression levels of a panel of centromere-associated genes. RT-qPCR was used to compare transcript levels in four independent PBRM1 knockout clones in 786-O, relative to transcript levels in parental cells. *GAPDH* and *PPIA* were used for normalisation. Points correspond to independent biological replicates. *n*=2-4, mean±SEM, and data were analysed by 2way ANOVA with Dunnett’s test, \**p*<0.05, \*\**p*<0.01, \*\*\**p*<0.001, \*\*\*\**p*<0.0001. **(c)** Clonogenic survival of 786-O parental and PBRM1 knockouts after *CCNB1* RNAi with two independent *CCNB1* siRNAs, normalised to survival after treatment with a scramble (scr) siRNA. Points correspond to independent biological replicates, *n*=4, mean±SEM, and data were analysed by 2way ANOVA with Dunnett’s test. **(d)** Representative image of colony formation from clonogenic survival assay, after treatment with the indicated siRNA as indicated in (c). **(e)&(g)** Clonogenic survival of 786-O parental and PBRM1 knockout cells after treatment with increasing doses of the Mps1 inhibitors (e) AZ3146, and (g) BOS172722, compared to survival with treatment with DMSO alone (Ø). *n*=4, mean±SEM, and data were analysed by 2way ANOVA with Dunnett’s test, \**p*<0.05. **(f)&(h)** Representative image of colony formation from clonogenic survival assays in (e) and (g).

**Figure S13.**
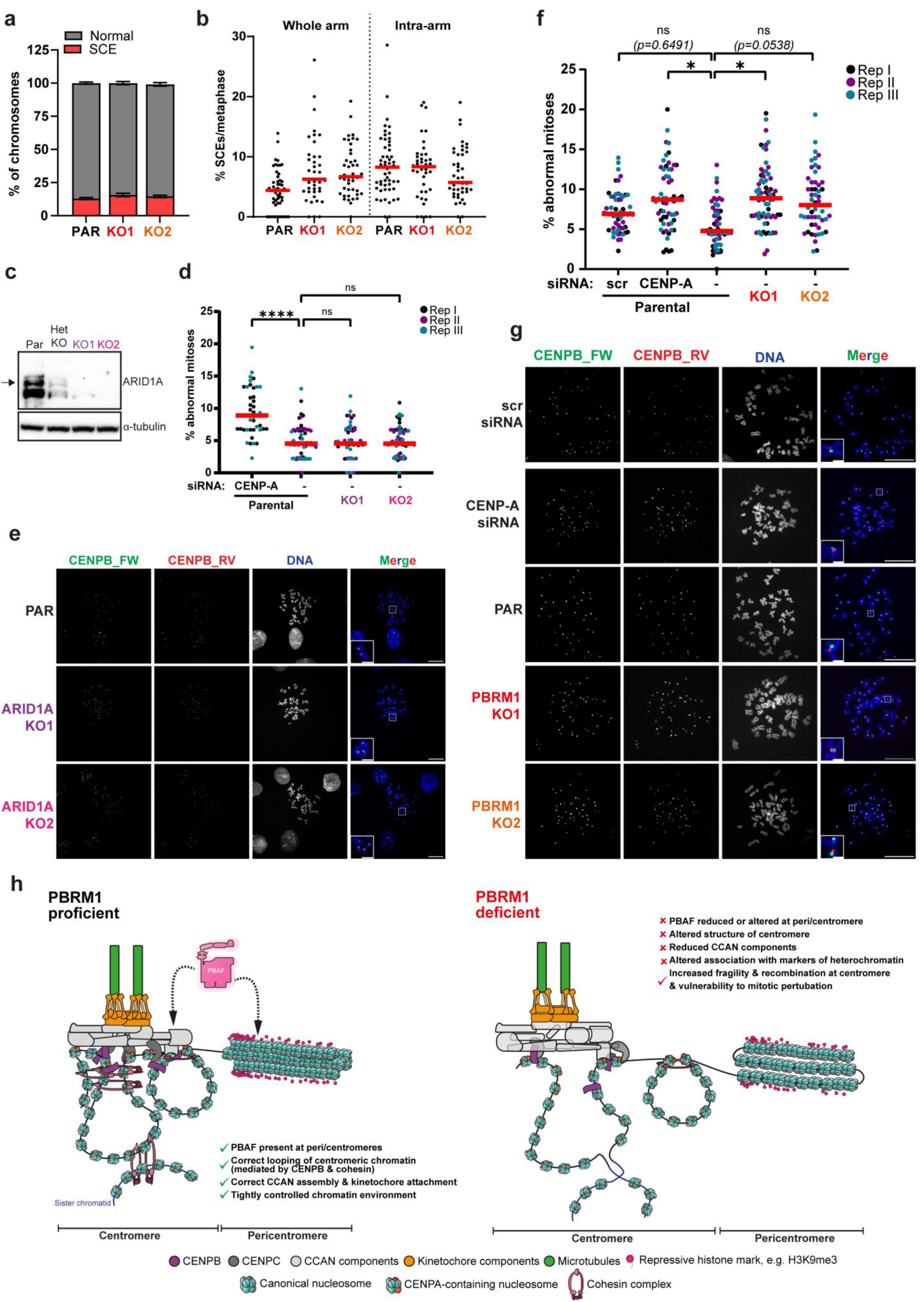
Centromere instability is conserved in other PBRM1 KOs but is not observed in ARID1A KOs. **(a)** Quantification of SCE assays from Figure 7(a)-(c), showing the percentage of chromosomes per metaphase which were either normal (grey) or had any SCE (red). **(b)** Quantification of the % of chromosomes in metaphases with either whole arm (left) or intra-arm (right) exchanges in RPE1 parental or PBRM1 knockouts. *n*=3, line at median. **(c)** Western blot for ARID1A in RPE1 parental and ARID1A knockouts (KO1 and KO2). A third clone (Het KO) was found to have a heterozygous mutation in ARID1A alleles and was not used in this study. α-tubulin is used as loading control, black arrow indicates band corresponding to ARID1A. **(d)** Quantification of chromatid exchanges at centromeres in RPE1 parental or ARID1A knockouts, or RPE1 parental cells, defined as the percentage of chromosomes with aberrant centromeres in each metaphase spread (% abnormal mitoses). *n*=3, line at median with different colours corresponding to independent biological replicates. In two replicates, RPE1 parental cells were also treated with CENP-A siRNAs to validate the detection of chromatid exchanges. At least 59 metaphases were analysed per condition (>40 in CENP-A condition), and data were analysed using 2way ANOVA with Dunnett’s test, \*\*\**p*<0.001. **(e)** Representative metaphase spreads from Cen-CO-FISH experiments in (d), with DNA stained with DAPI (blue), and centromeres hybridised with FISH probes against CENP-B-box sequences (green & red). Scale bars correspond to 20μm. White boxes correspond to zoomed inset images, where scale bars correspond to 5μm. **(f)** Quantification of chromatid exchanges at centromeres in 1BR3 parental or PBRM1 knockouts, or parental cells treated with CENP-A or scramble siRNAs, defined as the percentage of chromosomes with aberrant centromeres in each metaphase spread (% abnormal mitoses). *n*=3, with different colours corresponding to independent biological replicates, line at median, and data were analysed using 2way ANOVA with Dunnett’s test, \**p*<0.05. At least 50 metaphases were analysed per condition. **(g)** Representative metaphase spreads from Cen-CO-FISH experiments in (f), with DNA stained with DAPI (blue), and centromeres hybridised with FISH probes against CENPB-box sequences (green & red). Scale bars correspond to 20μm. White boxes correspond to zoomed inset images, where scale bars correspond to 5μm. **(h)** Model for how PBRM1 loss affect the centromeric and pericentromeric chromatin environment.

**Table S1:**
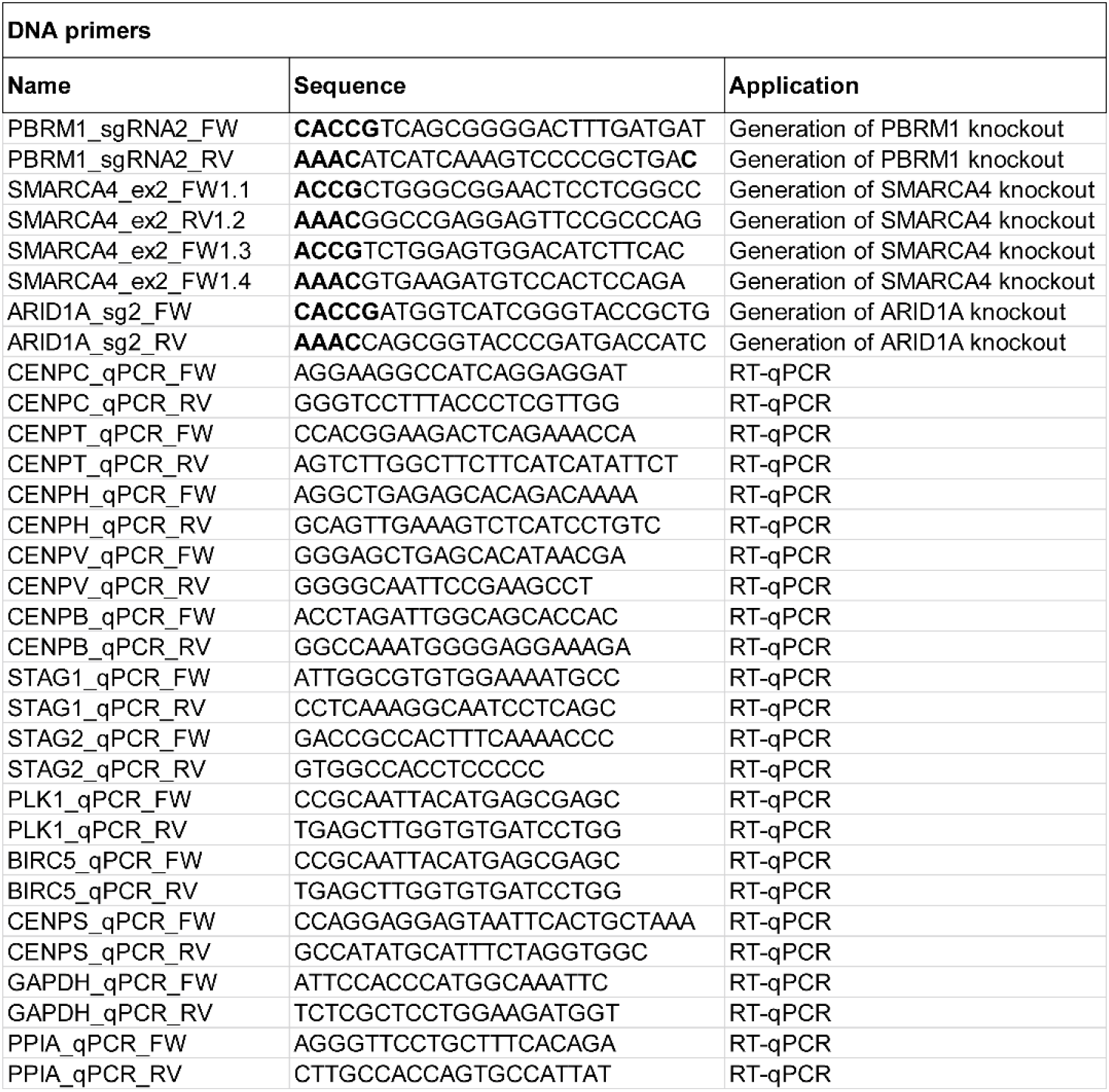
oligonucleotides used in this study.

**Table S2:**
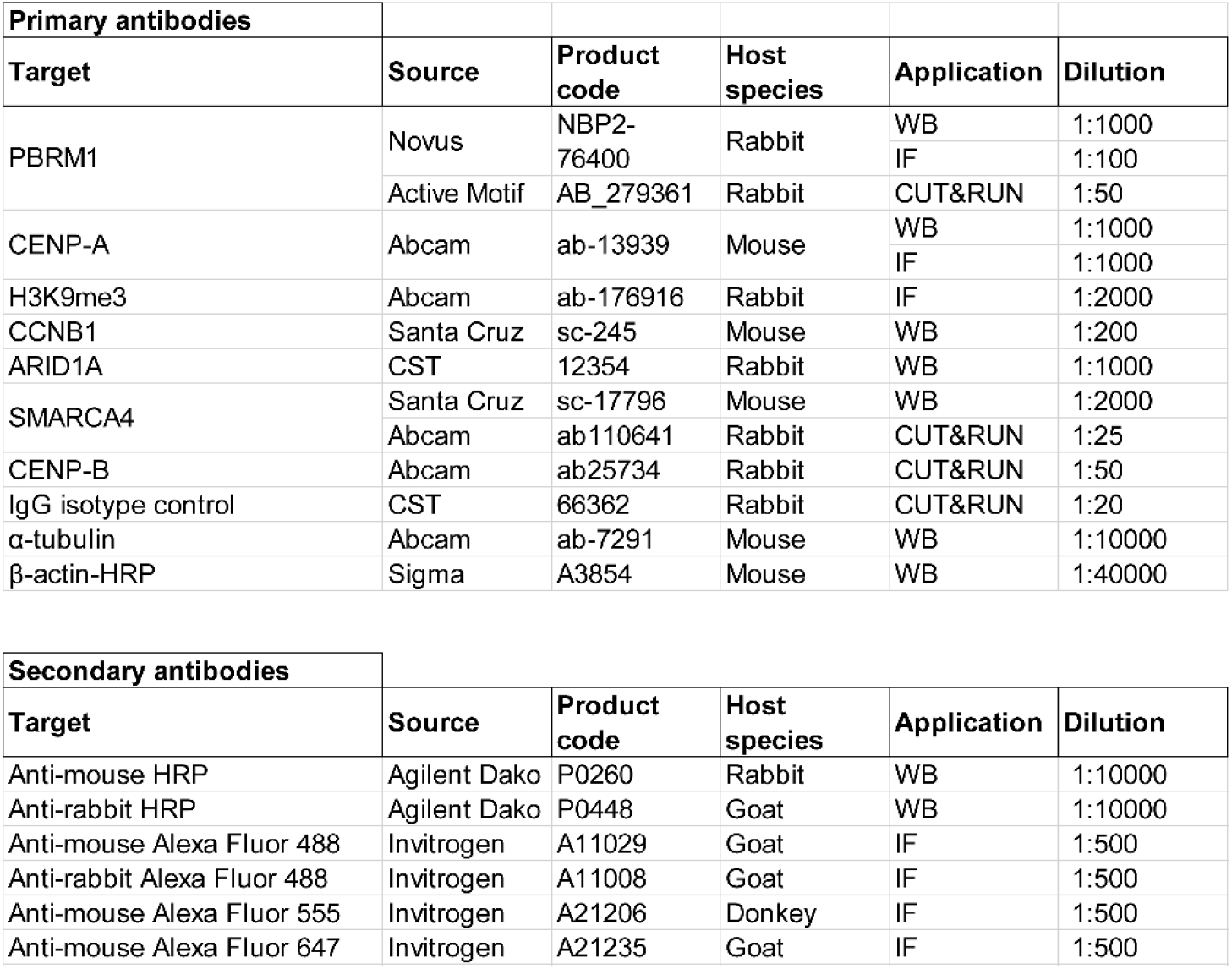
antibodies used in this study.

